# Oncogenic reactivation of young L1s is a hallmark of colon cancer

**DOI:** 10.1101/2023.05.17.541189

**Authors:** Devin Neu, Stevephen Hung, Cynthia F. Bartels, Zachary J. Faber, Katreya Lovrenert, W. Dean Pontius, Laura Morgan, Maharshi Chakraborty, Will Liao, Diana Chin, Ellen S. Hong, Jeremy Gray, Victor Moreno, Matthew Kalady, Ulrike Peters, Berkley Gryder, Richard C. Sallari, Peter C. Scacheri

## Abstract

Transposable elements become increasingly active in both cancerous and aging cells, driven by loss of DNA methylation as cells divide. Here we leverage the epigenomes of colon cancers with matched adjacent tissue, in addition to non-cancerous normals and cell line models, to assess the role of transposable elements as drivers or passengers in cancer development. Using the baseline of activity from normal and adjacent tissue, we show that the youngest subfamilies of the LINE1 (L1) family exhibit a degree of activity and recurrence across patients that goes beyond what is expected from hypomethylation and cell division, suggesting an additional mechanism of oncogenic reactivation. We characterize this mechanism and find that the loss of the tumor suppressor PLZF drives young L1 reactivation in a cell-division-independent manner. PLZF de-repression exposes abundant motifs for tumor core factors in the L1 5’UTR. Active young L1s act as oncogenic enhancers, interacting with oncogenes via gained chromatin loops. We uncover oncogenic L1 reactivation as a hallmark of colon cancer, where young L1s activate universally in our cohort at high levels of recurrence, act as enhancers to oncogenes, and become wired into the core regulatory circuitry of colon cancer.

## Main Text

Repetitive elements (REs) make up between one-half and two-thirds of the human genome. Most REs are derived from transposable elements (TEs), a small fraction of which are still autonomously mobile, namely the L1HS subfamily in the LINE-1 (L1) family^1, 2^. Rare, somatic L1HS insertions can disrupt the *APC* gene, a canonical tumor suppressor in colon cancer^3^. Non-mobile TEs can also become transcriptionally and epigenetically active in cancer cells, impacting genome function through their gene products or by acting as proximal and distal regulatory elements^4–6^. Additionally, TEs have been shown to act as alternative promoters for genes, and in some cases lead to activation of oncogenes such as *MET* in bladder cancer and *ARID3A* in lung cancer^7–9^. TEs are also active in aging cells. As cells divide, global hypomethylation increases driving TE reactivation^10–12^. In colon cancer, hypomethylation increases in the progression between normal, adenoma and carcinoma cells, continuing to raise gradually with tumor stage^13, 14^. Since TE reactivation is tied to the mitotic clock, increased activity is expected as tumor cells divide, confounding analyses that assess the role of TEs as drivers or passengers in cancer.

Here we interrogate the role of TEs in colon cancer while taking into account the expected increase in activity due to cell division. We leverage genome-wide maps of active and poised chromatin marks (H3K27ac, H3K4me1) in a cohort of 46 matched tumor and tumor-adjacent normal samples. We supplement the cohort with non-cancerous colon samples, which allow us to further interrogate the activity of TEs in colon tissue that is not poised to become cancerous. We also use a collection of model cell lines covering a range of stages in the progression from normal tissue to cancer. We find that the three youngest L1 subfamilies, L1HS, L1PA2 and L1PA3, exhibit a degree of activity and recurrence in tumors that is far greater than older L1s, suggesting an additional mechanism of L1 reactivation beyond cell division and hypomethylation. We show that these young L1s are oncogenic independently of retrotransposition, acting as enhancers that regulate oncogenes through gained chromatin loops. We characterize the mechanism driving the excess L1 activity and find that the loss of the tumor suppressor PLZF exposes abundant motifs for core regulatory factors of colon cancer in the 5’ UTRs of the young L1s. We term this “oncogenic TE reactivation”. In contrast to the rare instances where an L1 disrupts a tumor suppressor or oncogene through insertional mutagenesis, oncogenic TE reactivation of the three young L1 subfamilies is a hallmark feature of colon cancer.

## Results

### Young L1s enrich in recurrent, tumor-exclusive epigenetic activity

We performed ChIP-seq analysis of the active enhancer/promoter-associated histone modification, H3K27ac, across 46 colorectal tumor-normal pairs resected from human patients representing all clinical stages of disease (Supplementary Table 1). Each sample was sequenced to high depth (∼80M reads) using 150-bp paired-end reads; longer than typical for ChIP-seq. This facilitated characterization of regulatory elements at higher granularity than previous studies and enabled detection of TEs and other repetitive sequences in the genome with greater confidence.

We identify tumor-exclusive gains as H3K27ac peaks found in tumor samples that are absent in all other matched adjacent-normal samples. An unbiased survey against all simple repeats and major classes of repetitive REs comprising 56% of the human genome (Fig. 1a) reveals enrichment for three L1 subfamilies: L1HS, L1PA2, and L1PA3 (Fig. 1b). L1 sequences, like other REs, have expanded across the genomes of mammals over multiple rounds of mutation, retrotransposition and suppression by the host genome. Over the last 100 million years, dozens of L1 subfamilies have arisen from these cycles. L1MA subfamilies are found across all mammals, the L1PAs in primates, and the L1HS subfamily is exclusive to *Homo sapiens*. We find 79% of the young L1s that overlap tumor-exclusive gains are full-length (fl-L1s), five-fold greater than expected (Supplementary Table 2). Furthermore, H3K27ac signal at L1s is concentrated within the first 200 bp of their 5’UTR (Fig. 1e,f). While repetitive elements have traditionally complicated read-mapping, we verify that we can reliably identify H3K27ac signal at L1s using paired-end 150– bp reads aligned to non-repetitive flanking regions, with ChIP input uniformly covering the rest of the L1 body (Extended Data Fig. 1). The fl-L1 gains also exhibited increased transcription, further supporting their gain of activity in tumors (Extended Data Fig. 2). We establish fl-L1 5’UTRs, among all RE classes, as our primary focus to investigate the role of TEs in the transition between normal and cancer cells in colon.

**Fig. 1.**
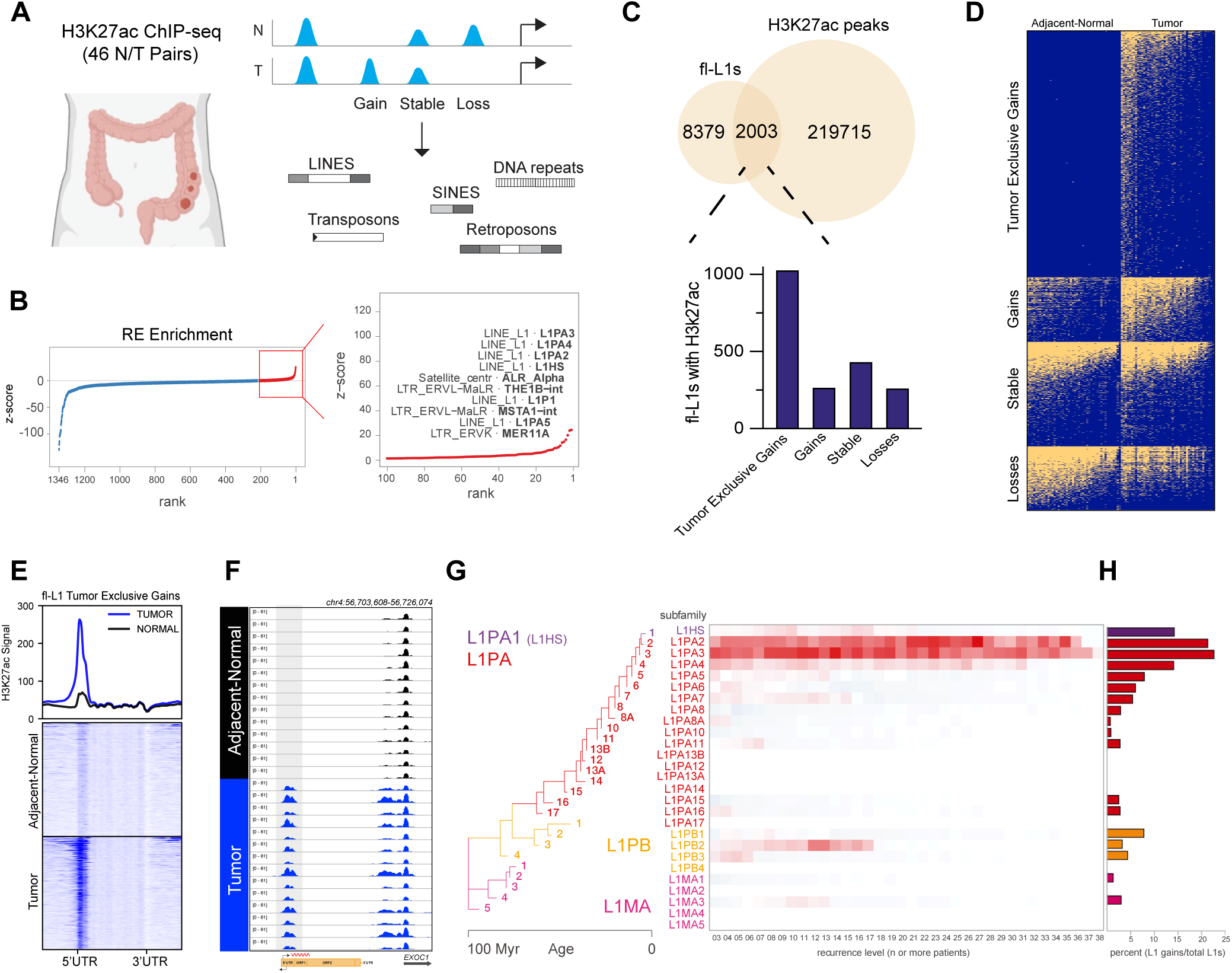
Young L1s enrich in recurrent, tumor-exclusive epigenetic activity. **a**, Schematic of experimental strategy. Tumor specific gains and losses of H3K27ac were identified and intersected with all classes of repetitive elements (RE). **b**, Enrichment of all repetitive element classes at tumor specific H3K27ac gains with ranked z-scores. Top 100 most enriched RE classes are marked in red and highlighted to the right. **c**, Intersection of all full-length L1s with H3K27ac sites in adjacent-normal and tumor with categorical assignment based on status within the cohort. **d**, Binary heatmap of fl-L1 activity status across the full cohort, on (yellow) and off (blue). **e**, Aggregate H3K27ac signal for all 46 adjacent-normals (black) and tumors (blue) at tumor– exclusive L1 gains. **f**, Representative browser tracks of a recurrently reactivated L1 with H3K27ac ChIP-seq for adjacent-normals (black) and tumors (blue) at *EXOC1* locus with L1 reactivation displayed upstream highlighted in gray. **g**, Phylogenetic tree of L1 evolution over 100 million years, heatmap displays enrichment z-scores for L1 overlap with H3K27ac elements of increasing recurrence level in patients (x-axis). **h**, Percent of each L1 subfamily gains in relation to the total number of L1s in each subfamily (reactivation rates).

We retrieved 8,319 fl-L1 5’UTRs in the human genome and determined their overlap with the union of all H3K27ac peaks in our cohort (219,715 non-overlapping). 2,003 fl-L1s 5’ UTRs overlapped H3K27ac peaks in two or more of the 96 samples. Based on their activity in adjacent-normal and tumor samples we group them into gains, losses or stable L1s. We further subdivide gains into a group of tumor-exclusive gains, with no activity across adjacent-normal samples. Half of the fl-L1 5’UTRs exhibit activity in adjacent-normal tissue while the other half (51%) are exclusive to tumor (Fig. 1c,d). The balance of L1 gain and loss is highly asymmetrical; only 0.02% of fl-L1 5’UTRs are exclusive to adjacent-normal. On average, H3K27ac signal at tumor-exclusive fl-L1 5’ UTRs is 6.7-fold higher in tumor relative to adjacent-normal (Fig. 1e).

Active fl-L1s showed striking levels of recurrence, with individual L1s gaining H3K27ac signal in nearly every tumor evaluated (Fig. 1f). Tumor-exclusive recurrent H3K27ac regions, active in a high number of tumor tissues, overlap young fl-L1s, L1PA2 and L1PA3, with surprising frequency. Of 236 H3K27ac gains recurrent in at least half of the tumor tissues (23 or more out of 46), 31 (13%) overlap young fl-L1s (Supplementary Table 3). Overlaps increase with the level of recurrence of active fl-L1s, reaching 25% (9/36) at two-thirds or more of all tumors. Among all L1 subfamilies across all recurrence levels, L1PA2 and L1PA3 exhibit a stark enrichment when compared to a null model obtained from overlapping L1s with permuted subsets of the tumor-exclusive H3K27ac gains, with the strongest signal for L1PA3s at 12/46 recurrence (-log_10_(*P*)=13.85, (Fig. 1g, See Methods). Average reactivation rates in the cohort are highly variable across L1 subfamilies, ranging from 0% in old and some middle-aged subfamilies to over 22% for L1PA2s and L1PA3s (Fig. 1h). Generally, L1 reactivation rates decrease with evolutionary age.

### Young L1s have greater activity and recurrence than expected

Having established that L1s are important in tumor exclusive epigenetic activity, we set out to quantify the rate at which they become active by gaining H3K27ac, and the extent to which this rate differs between non-cancerous normal, tumor-adjacent normal, and tumor. We calculated the reactivation rate of each L1 subfamily by comparing the number of L1s that gain H3K27ac and dividing by the total number of L1s in that subfamily.

We established a baseline of L1 reactivation by profiling non-cancerous normal colon tissue, since adjacent-normal tissue may already harbor some of the epigenetic alterations we observe in the tumor tissue. Generally, reactivation rates don’t exceed one or two percentage points (mean: 1.3%) in non-cancerous colon tissue. The tumor-exclusive young L1 subfamilies were especially muted, however two patients broke the trend, reaching 6.7% and 7.1% in the L1PA2s (Fig. 2a). In adjacent-normal, L1PA2s showed the highest reactivation rates (mean: 10.4%, SD: 3.5%, max: 15.5%) and were active in all but two samples. We did not observe a general trend with evolutionary age; even relatively young families, like L1PA5s, remain inactive (mean: 1.2%, SD: 0.7%, max: 3.0%). Subfamily reactivation rates in adjacent-normal samples showed remarkable uniformity, with an average SD of 1.3%, resulting in a striped pattern in the heatmaps (Fig. 2a). Tissues collected from patients from separate countries (US and Spain) are indistinguishable.

**Fig. 2.**
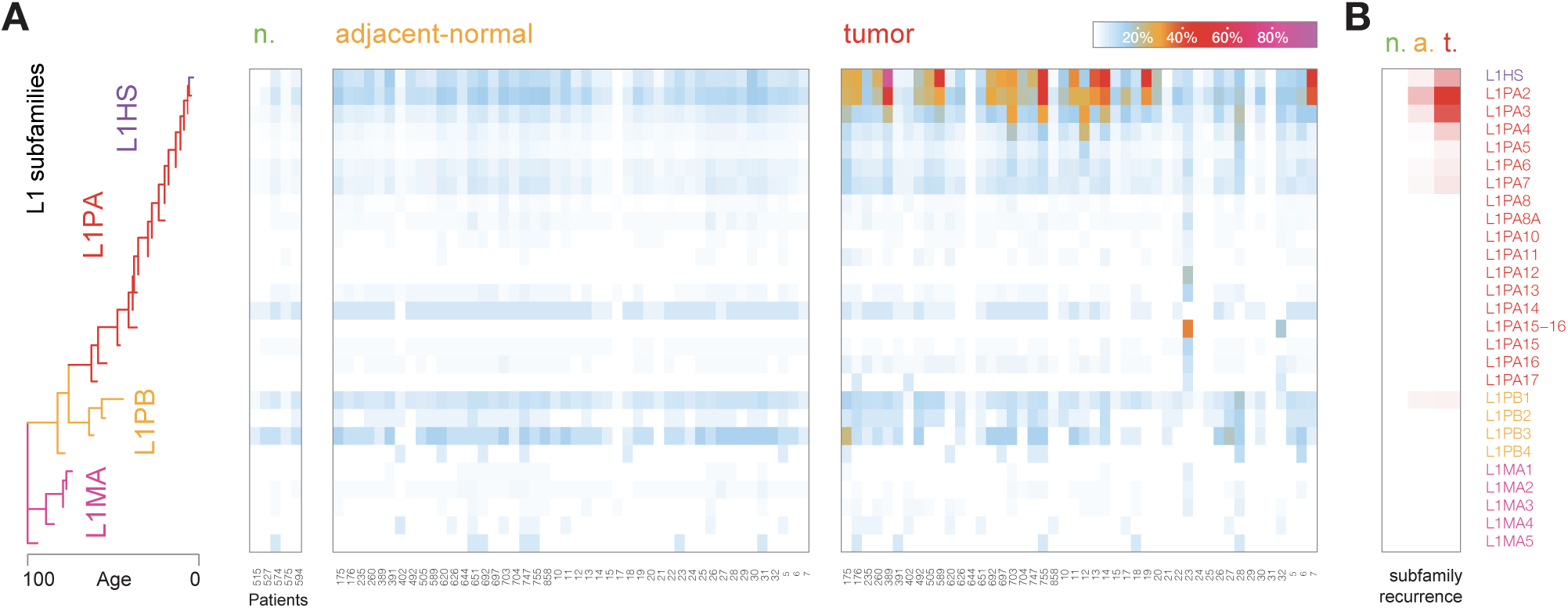
Young L1s have greater activity and recurrence than expected. **a**, L1 phylogenetic tree, heatmap displays percentage of reactivated L1s for each subfamily (rows) across non-cancerous-normal, adjacent-normal and tumor samples (columns). **b**, Heatmap showing the degree of subfamily recurrence across L1 subfamilies (rows) and cancer stages (columns) from low (white) to high (red).

In tumors, reactivation rates were similar for most subfamilies and exhibited the familiar striped pattern found in adjacent-normal with the notable exception of the four young subfamilies, L1HS, L1PA2, 3 and 4, whose reactivation rates increased 2.4-fold on average. Reactivation rates across patients also became highly variable, with L1HSs (mean: 19.0%, SD: 17.3%, max: 85.2%) and L1PA2s (mean: 18.7%, SD: 11.1%, max: 50.4%) reaching staggering levels of activity (Fig. 2a). The older L1 subfamilies showed more modest and uniform increases. We interrogated the difference between young and old L1s by comparing their increase in activity across tumor T-stage. L1 hypomethylation increases linearly across adjacent-normal, adenoma and the four T-stages of colon cancer (Extended Data Fig. 3a), where each transition sees a ∼10% increase in activity ^13^. The older L1PA subfamilies showed a gradual increase in H3K27ac activity that tracked with the hypomethylation data. However, the three youngest L1 subfamilies showed a marked deviation from the trend that suggested an additional mechanism beyond what could be explained by hypomethylation (Extended Data Fig. 3b). The excess levels of activity in the youngest L1 subfamilies did not appear to be driven by increased H3K27ac activity in their flanking regions (Extended Data Fig. 3c, see Methods).

We next assessed the degree of recurrence across patients for each L1 subfamily in each of the three cell states: non-cancerous-normal, adjacent-normal and tumor. The high reactivation rates we observe in the young L1 subfamilies do not necessarily imply high levels of recurrence for individual L1 elements across patients. We designed a subfamily test of recurrence based on a random model of L1 reactivation (Extended Data Fig. 3d, see Methods). Under the random model, the L1PA2 subfamily, with an average reactivation rate in tumor samples of 18.7%, we expected the majority of L1s to be active in five to 16 patients, and virtually none in over 33. We observed 82 L1PA2s active in more than 33 tumor samples (-log_10_(*P*)=65.21). We detected significant recurrence in the three youngest L1 subfamilies in both adjacent-tumor and tumor states, and in the L1PA4, 6, 7 and L1PB1 subfamilies in the tumor state alone (Fig. 2b). We found no evidence of recurrence in the non-cancerous-normals. Our findings suggest that excess activity in young L1s may be an early event in colon cancer. Further evolutionary structure in L1PA2s did not account for recurrence (Extended Data Fig. 4a,b, see Methods). In summary, L1 reactivation is universal in our cohort, with the youngest L1 subfamilies exhibiting exceptionally high activity and recurrence in tumor. By contrast, older L1 subfamilies with comparable activity in adjacent-normal tissue appear to conform to a gradual increase consistent with expanding hypomethylation driven by cell division.

### L1s act as oncogenic enhancers

We performed a QTL-like analysis in which we identified patients with or without L1 reactivation, and then compared the expression of L1 target genes based on reactivation status. If fl-L1s function as enhancers, we expect to see increased expression of L1 gene targets in patients that have a reactivated L1. Overall, 22% of the total set of reactivated L1s correlated with the expression of a gene. Consistent with their role as enhancers, 80% of the top 100 gene targets showed increased expression when the L1 peak was present with a median fold change of 2 (Extended Data Fig. 5a-d).

We performed H3K27ac HiChIP analysis of an adjacent-normal and tumor pair (patient 28) to test if reactivated L1s loop to overexpressed genes. We found that 797 reactivated L1s form new loops to 18 overexpressed genes in our cohort (Extended Data Fig. 5e-g). We identified 457 reactivated L1s which share topologically associated domains (TADs) with 70 known oncogenes (Fig. 3a). In general, the contact frequencies between these L1s and the oncogenes within TADs increased from normal to tumor (68.8% increase, p-value: 0.038). Among 87 possible connections, 10 showed either a completely new loop or an increased contact frequency between the L1 and the oncogene in the tumor sample relative to its matched adjacent-normal control (Fig. 3a). These oncogenes and their corresponding fold changes in contact frequency are as follows: *BCL11A* (14.25X), *KLF4* (4.92X), *MMS22L* (4.56X), *COPS5* (4.50X), *YWHAZ* (3.82X), *BCL6* (2.50X), *HSPA4* (2.00X), *TFG* (1.77X), *FGFR1OP* (1.40X), and *ETV6* (1.10X). Reactivated L1s in patient 28 also showed new loop interactions outside of oncogenes, including hundreds of interactions to tumor-specific H3K27ac gains (Fig. 3b). The L1 interaction with *BCL11A*, an oncogene involved in lymphoma, lung and breast cancer where it activates Wnt/β-catenin signaling, showed a switch-like behavior with the oncogene promoter (Fig. 3c). In adjacent-normal the TSS interacts with a pair of proximal enhancers, whereas in tumors adjacent-normal loops are lost and replaced by two novel, punctate loops connecting the oncogene promoter directly with the L1’s 5’ UTR and a second neighboring enhancer. The gained *BCL11A* loop is a clear example where the L1 dominates the oncogene’s regulatory landscape and appears to erase all previous chromatin structure. The L1 interaction with *KLF4*, one of the four core iPSC reprogramming transcription factors, exemplifies an L1 behaving additively by joining a collection of 7-9 gained loops within the gene’s TAD (Fig. 3d). Both examples of contacts between L1 5’UTRs and the *BCL11A* and *KLF4* promoters underscore the specificity and strength of gained loops to oncogenes in tumors. Finally, recurrent L1s were four times more connected within their TADs when compared to their reactivated but non-recurrent counterparts (Fig. 3e,f, –log_10_(*P*)=18.791).

**Fig. 3.**
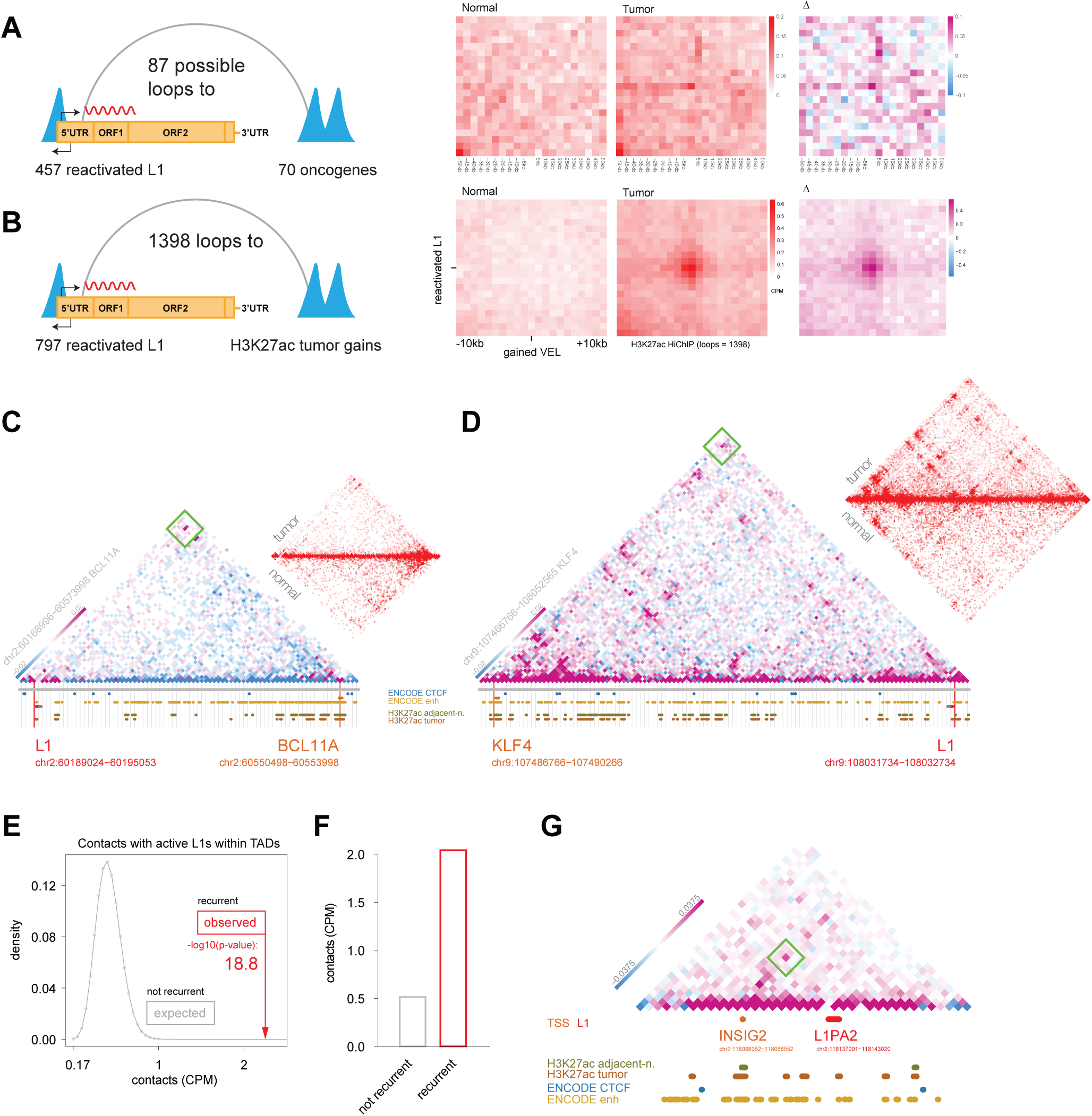
L1s act as oncogenic enhancers. **a**, (Left) Diagram of L1s to oncogene loops. (Right) Aggregate Peak Analysis (APA) plots of H3K27ac HiChIP. Aggregate contact frequencies of adjacent-normal (left), tumor (middle), and the delta (tumor minus normal) (right). **b**, Same as a, but for L1 loops to H3K27ac tumor gains. **c**, L1 loop gain to BCL11A oncogene. The contact map (5 kb resolution) represents the delta between tumor and normal, gains (purple) loss (blue). The gained loop peak between the L1 and BCL11A is framed in green. The raw contact frequencies are shown in the upper right red-white heatmap for tumor (top) and normal (bottom). **d**, L1 loop gain to KLF4 **e**, Statistical analysis comparing observed contacts between recurrent L1s and neighboring regulatory elements (vertical red arrow) compared to a null distribution obtained from non-recurrent L1s (gray curve). **f**, Observed average contact frequencies for recurrent L1s compared to expected by the null model. **g**, Indicated in gray is a highly recurrent, pro-growth enhancer of the L1PA2 subfamily. The region exhibits no conformational structure in the normal-adjacent sample but gains a loop in the tumor between the L1 and the INSIG2 gene TSS.

A pro-growth enhancer identified from a CRISPRi screen in colon cancer cells^15^ overlapped a recurrent L1 active in patient 28. Here we show that the oncogenic L1 establishes a novel interaction with *INSIG2* (Fig. 3g), a gene reported to have prognostic capacity for colon cancer survivorship; its overexpression in cancer cells results in numerous cell phenotypes related to growth, invasion and apoptosis, appearing to suppress the efficacy of chemotherapy^15, 16^. Overall, these findings indicate that reactivated young L1s can act as enhancers, altering gene expression. Furthermore, L1s can directly target oncogenes through highly punctate and tumor-specific chromatin loops. Finally, silencing the enhancer activity of a single, L1 can have a major impact on tumor cell phenotype indicating that L1s can possess intrinsic oncogenicity.

### L1 gains are active in stem cells and poised for reactivation in adjacent-normal colon

In humans, L1s are active in embryonic tissues and cells and later become silenced during differentiation, while some persist or are re-expressed during lineage specification^17–19^. This prompted us to evaluate if the active L1s in colon tumors are active in normal tissues or at the early stages of development, as this could provide insights into their origin. The Roadmap project surveyed the tissue specificity of promoters and enhancers, providing modules that group regulatory elements based on their activity patterns across 111 human tissues. Roadmap modules range in tissue specificity from constitutive, or active in virtually every cell type, to highly specific, capturing the epigenetic identity of an individual cell type or lineage. Compared to direct overlap with cell-type annotation, which contain a mix of constitutive and specific elements, this approach provides a direct measure of how broadly L1 subfamilies activate across human tissues. We overlapped CRC-active fl-L1s with Roadmap modules by subfamily (Fig. 4a). One-third of active fl-L1s in colon tumors overlapped Roadmap modules. The highest overlap was between the PA2 and PA3 subfamilies and two stem-cell-specific Roadmap promoter modules: pro41and pro74, comprised the majority (58.76%) of the Roadmap-overlapping fl-L1s from the young subfamilies (Fig. 4b). Module pro41 is primarily active in both embryonic stem cells and induced pluripotent stem cells (iPSCs), while module pro74 is primarily active in iPSCs. L1 subfamilies older than L1PA6 notably lack these two specific modules, suggesting that activity in an embryonic state is a distinct feature of younger L1 subfamilies.

**Fig. 4.**
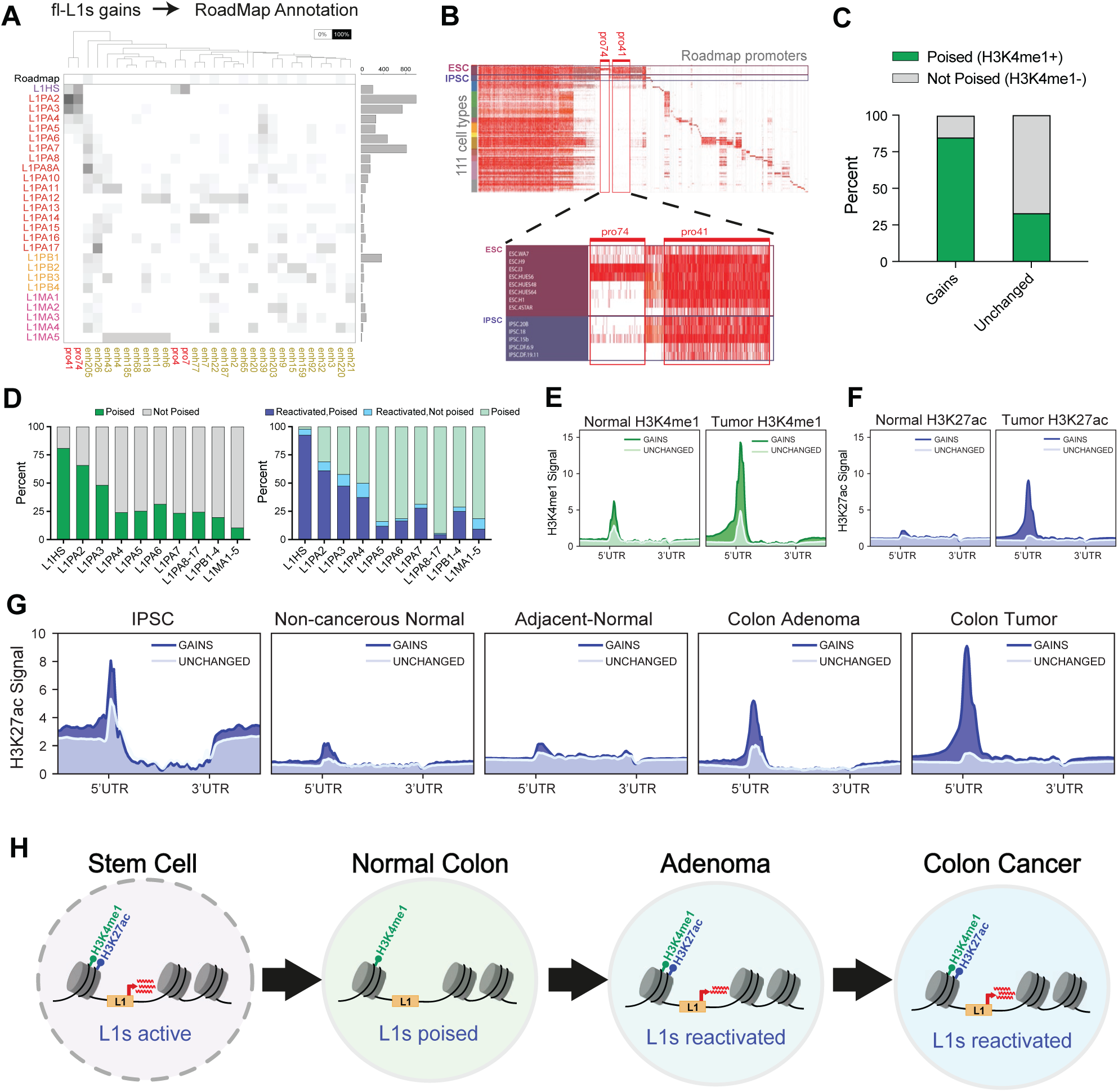
L1 gains are active in stem cells and poised for reactivation in adjacent-normal colon. **a**, Percent overlap between fl-L1s gains and Roadmap promoter (red labels) and enhancer (gold labels) activity modules. The designated names of the modules are on the x-axis, while the y-axis lists L1 family names, sorted by evolutionary age. Bar plot designating L1 counts (right). **b**, Roadmap promoter modules across 111 normal cell types (y-axis) and all putative human promoters (x-axis, adapted from Roadmap). Modules pro74 and pro41 are highlighted and blown up to show the specificity of their activity in ESCs and IPSCs. **c**, H3K4me1 ChIP-seq (CRC-14) peaks intersected with sample specific H3K27ac gains to calculate the percent of L1 gains poised in adjacent normal. **d**, Percent of L1 subfamilies poised in normal (green), percent poised L1s that become reactivated in tumor (blue), percent of L1s not poised in normal but are reactivated in tumor (light blue). **e**, H3K4me1 and **f**, H3K27ac ChIP-seq signal at gains compared to unchanged fl-L1s for CRC-14. **g**, H3K27ac signal at gains and unchanged fl-L1s across different stages and cell types illustrating developmental progression of L1 reactivation. **h**, Model of L1 reactivation through colon cancer tumor formation.

For two paired samples (CRC-14 & CRC-28), H3K4me1 ChIP-seq was performed to evaluate the poised status of the fl-L1 gains in both adjacent-normal and tumor. After intersecting H3K4me1 poised peaks with CRC-14 and CRC-28 specific gains, we found that 84% and 59% of fl-L1 gains in those samples are poised in their matched adjacent-normal, respectively (Fig. 4c, Extended Data Fig. 6a). In comparison, 33% and 21% of unchanged fl-L1 (those lacking H3K27ac signal in either adjacent-normal or tumor) were poised in adjacent-normal. (Fig. 4c, Extended Data Fig. 6a). Given that young L1 subfamilies have high reactivation rates, we also evaluated the poised status within each L1 subfamily. For simplicity of analyses, older subfamilies were combined into 3 groups past L1PA7 (L1PA8-17, L1PB1-4, and L1MA1-5). We find that the percent of tumor gains poised in adjacent-normal were highest in young L1 subfamilies and decline with evolutionary age (Fig. 4d, Extended Data Fig. 6b). Next, we determined the percentage of these poised fl-L1s in adjacent-normal that become reactivated in tumors. Once again, younger subfamilies showed higher reactivation rates and we saw a decay in reactivation with evolutionary age. Importantly, there are only rare instances where L1s became reactivated in the tumor that were not poised in adjacent normal, demonstrating poised status is a key predictor of L1 reactivation. Together, young L1 subfamilies are more poised for reactivation, and subsequently have higher reactivation rates in tumors. In looking at the H3K4me1 ChIP-seq signal, we find that both gains and unchanged fl-L1s are poised to similar levels in adjacent-normal, with gains showing slightly higher signal (Fig. 4e, Extended Data Fig. 6c). In tumor fl-L1 gains show increased H3K4me1 signal compared to unchanged fl-L1s in addition to showing specific H3K27ac gain in tumor (Fig. 4f, Extended Data Fig. 6c).

We next investigated the timing of L1 reactivation in the development of colon cancer. In addition to the adjacent-normal samples for each pair, we profiled two colon adenoma samples and 5 normal colon samples resected from non-cancerous patients, or non-cancerous normals. Using sample 14’s gains and unchanged fl-L1s, we find these sites are active in an iPSC cell line further supporting the Roadmap result. When looking at both non-cancerous normal and adjacent normal we see little to no H3K27ac signal. Interestingly, we do observe H3K27ac reactivation begin in the adenoma suggesting L1 reactivation occurs during the early stages of tumor formation and continues to increase in colon cancer (Fig. 4g, Extended Data Fig. 6d). Taken together a model emerges where young L1s are active in embryonic cell states, largely become silenced in lineage-defined tissue and then reactivate at increasingly high rates during tumor formation (Fig. 4h). Cells become increasingly undifferentiated and adopt a more stem-like state during tumorigenesis, which could explain why we observe specific reactivation of young fl-L1s as they are specifically enriched for active stem cell signatures and factors. Whether the reactivation of young fl-L1s is secondary to the cells becoming increasingly stem-like or an actual determinant of the stem-like state remains an open question.

### L1s are wired into the core regulatory circuitry of colon cancer

We identified the core-regulatory circuitry of normal colon and colon cancer based on super-enhancer profiles and expression^20, 21^. We uncovered distinct circuitries for both adjacent-normal and tumor in addition to a shared circuitry (Fig. 5a, Extended Data Fig. 7a). For example CREB1 is a core-regulatory factor (CR TF) in the tumor circuitry absent from the normal core-regulatory circuitry, while VDR is a shared factor, and HIC1 is a normal-specific core factor. An unbiased motif search of the reactivated L1s 5’UTR revealed an overabundance of these and many other CR TF motifs, where they rank among the most highly significant of all motifs detected (Fig. 5a). Additionally, digestive-specific TF ChIP-seqs (882 total) were used to intersect the H3K27ac sites to corroborate motif mining data in a lineage context (Extended Data Fig. 7b). Upon further inspection of the motifs with respect to the 5’UTR, we found that they concentrate within the H3K27ac peak and in some cases multiple instances of the core-regulatory motifs are observed (Extended Data Fig. 7c).

**Fig. 5.**
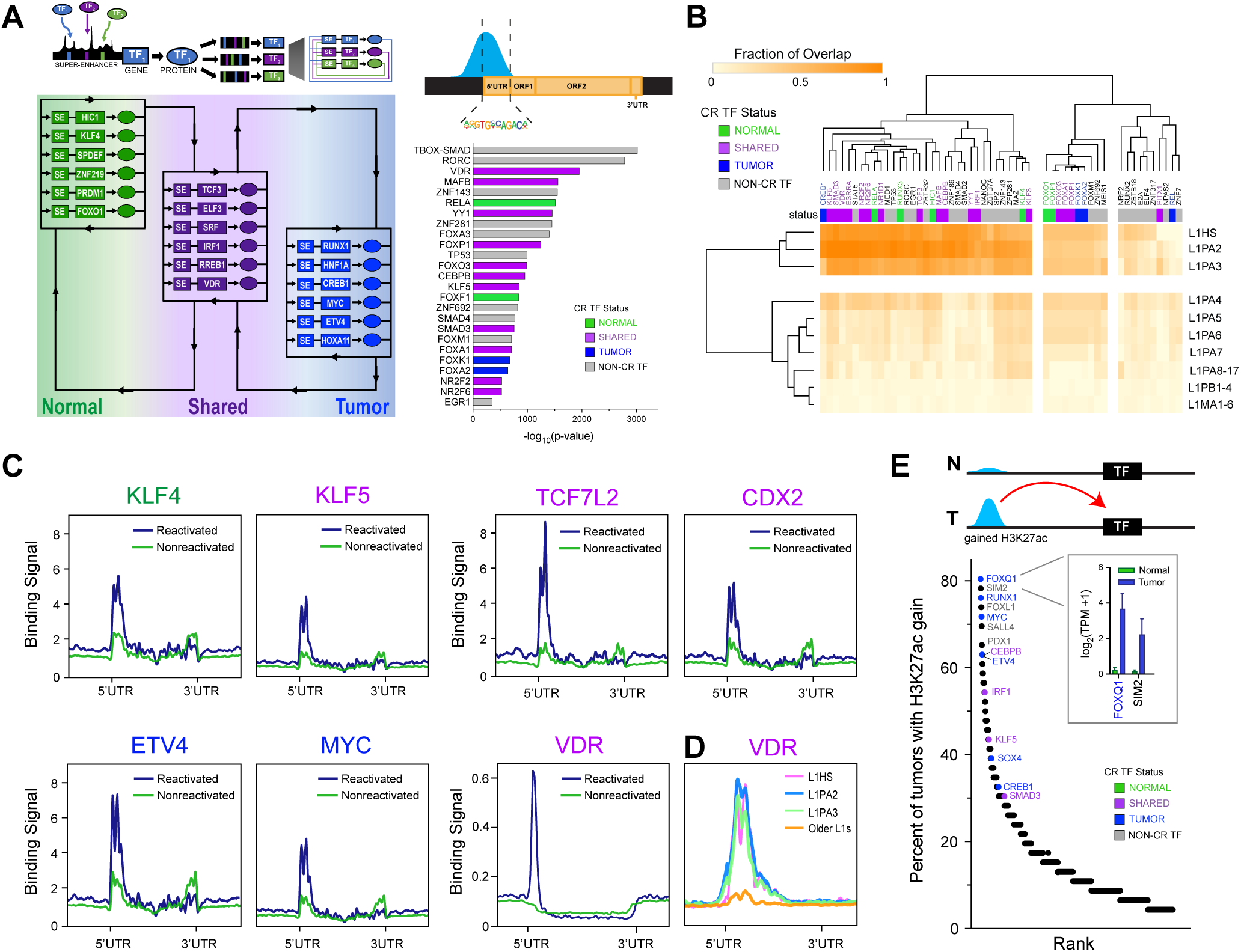
L1s are wired into the core regulatory circuitry of colon cancer. **a**, Model of core regulatory circuitries identified in adjacent-normal and tumor, top 6 ranked CR TFs for each category based on degree of connectivity are shown, motifs for many of these CR TFs are enriched at reactivated L1s. **b**, Unsupervised clustering of the fraction of motif overlap for L1 subfamilies, older subfamilies are combined for L1PA8-17, L1PB1-4, and L1MA1-6. CR TFs labeled according to circuitry status. **c**, TF binding profiles for CR TFs at the L1 5’UTR, CUT&RUN TFs (KLF4, KLF5, TCF7L2, CDX2, ETV4, and MYC); ChIP-seq (VDR, publicly available) with binding signal plotted at reactivated fl-L1s (blue) compared to non-reactivated fl-L1s (green). **d**, VDR binding signal at the 3 youngest subfamilies and combined older L1 subfamilies at the 5’UTR. **e**, Percent of patients with gained H3K27ac sites in tumors associated with TFs ranked by recurrence.

To assess whether the CR TFs could be accounting for the exceptionally high reactivation rates in the young L1 subfamilies, we evaluated the subfamily motif composition across evolutionary time and clustered the enriched motifs at the subfamily level based on the fraction of overlap for those L1 sites (Fig. 5b). The three youngest L1 subfamilies, L1HS, L1PA2 and L1PA3, contained an excess of motifs for a large group of core regulatory factors. This excess is especially pronounced in the separation between L1PA4 and L1PA3 where unbiased clustering of motif abundance starkly separates the young from the older L1 subfamilies. We verified that several of these core factors bind the reactivated L1s using CUT&RUN (KLF4, KLF5, TCF7L2, CDX2, ETV4, MYC) and publicly available ChIP-seq data (VDR) (Fig. 5c). Furthermore these core factors showed higher enrichment at the young L1s than the older L1s (Fig. 5d, Extended Data Fig. 7d). Core factor motifs are enriched in the consensus sequences of all young L1s, but only a fraction recur across patients. We further compared core factor motif composition between highly recurrent and completely inactive instances within the L1PA2s (Extended Data Fig. 8a,b). Recurrent L1PA2s contain an excess of core factor motifs, including MYC, KLF4/5 and NANOG, that result from L1-locus-specific sequence variants.

With the exception of *APC* mutated in 80% of colorectal tumors, no other known CRC driver reaches a somatic mutation rate that alone is likely to account for the universality of L1 reactivation we observe. We reasoned that an epigenetic driver event is more likely, as we and others have shown that the frequency of recurrent epigenetic events, especially at enhancer elements, is often greater than recurrent mutational events^22^. In search of such a high-frequency epigenetic event, we took all H3K27ac gains and losses in tumors relative to each of their matched controls, identified those that target transcription factors, and ranked them by their level of recurrence. Gains targeting CR TFs including FOXQ1, RUNX1, MYC, CEBPB, ETV4, and KLF5 are among the most recurrently gained enhancers (Fig. 5e). Core factors bind reactivated L1 elements, but the observation that poised elements with same motifs are not bound by the core factors indicates that an additional factor is needed for L1 reactivation. We hypothesized this factor is likely to be a repressor of L1 that is lost during tumorigenesis, leading to an unmasking of the core regulatory motifs and enabling binding of the core-regulatory TFs. By virtue of both being regulated by core factors and in turn regulating oncogenes and other core factors, L1s are wired into the core regulatory circuitry of colon cancer.

### PLZF drives oncogenic reactivation of young L1s

As noted above, despite the CR TFs being available and the L1s being poised in adjacent-normal, we hypothesized that some additional factor is necessary to fully activate L1s. We homed in on *ZBTB16*, a tumor suppressor in many cancers and L1 repressor in myeloid progenitor cells^23–26^. Although rarely mutated in colon cancer, we find *ZBTB16* (PLZF) is frequently dysregulated at the epigenetic level ranking 5th among the TFs associated with recurrent H3K27ac losses (Fig. 6a). The magnitude of the loss at the PLZF locus, i.e. the difference in H3K27ac signal (RPKM) between adjacent-normal and tumor has a 60% decrease (q-value<1e10^5^) (Fig. 6a-c). This epigenetic loss of PLZF activity is reflected transcriptionally, with significant downregulation in human tumors (Fig. 6d).

**Fig. 6.**
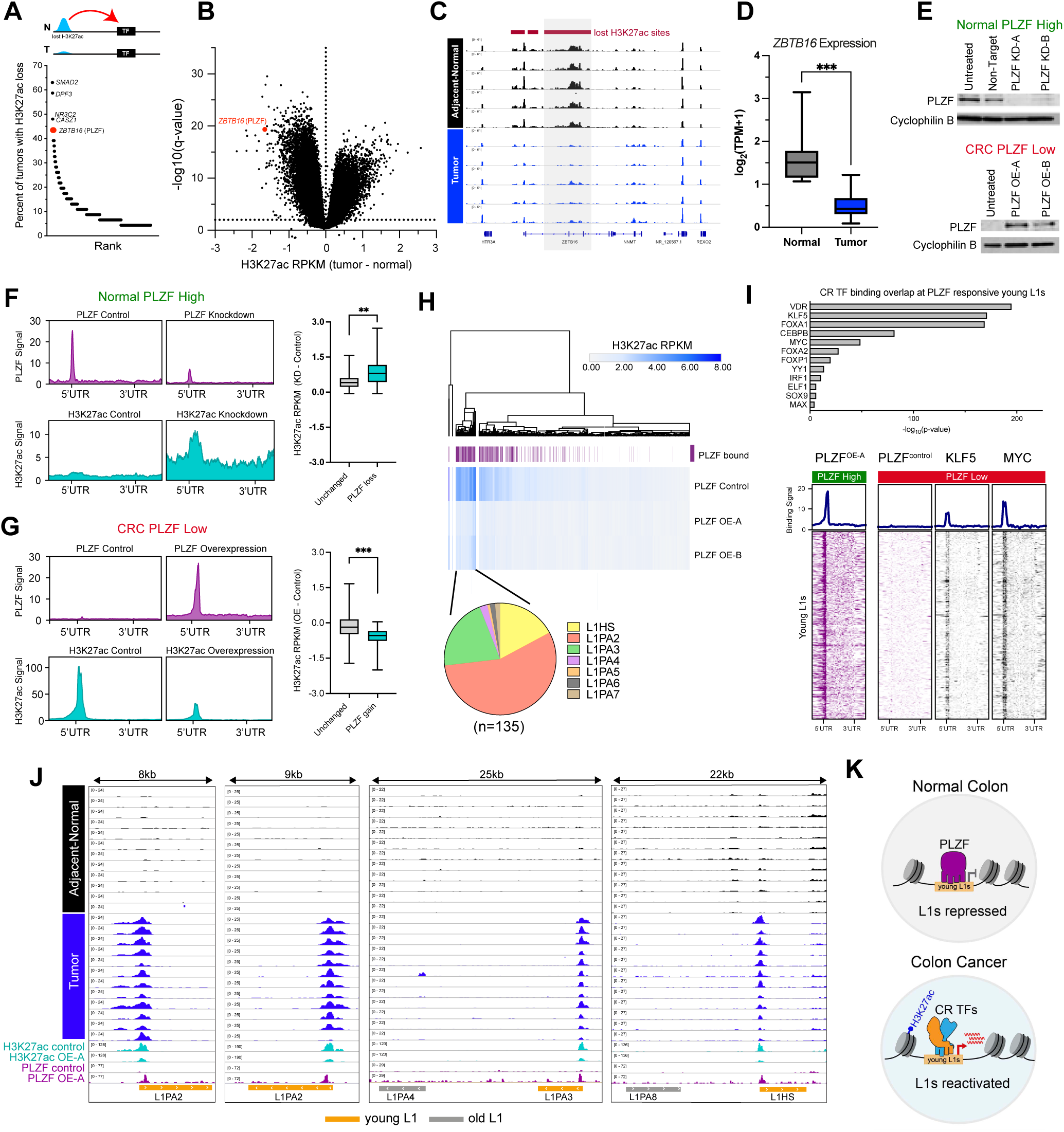
PLZF drives oncogenic reactivation of young L1s. **a**, Percent of patients with lost H3K27ac sites associated with TFs ranked by recurrence. Top 5 lost H3K27ac associated TFs labeled **b**, Volcano plot of H3K27ac RPKM differences between adjacent-normal and tumor (log_10_q-value). ZBTB16 lost H3K27ac sites labeled in red. **c**, Representative H3K27ac tracks at ZBTB16 locus of lost H3K27ac signal between adjacent normal and tumors, recurrent H3K27ac loss associated with ZBTB16 boxed in gray. **d**, log_2_TPM gene expression of ZBTB16 in adjacent normal and tumor (*** p<0.001) **e**, Western blot of PLZF for knockdown in normal (FHC) colon line, untreated, and non-target siRNA controls. PLZF overexpression in CRC cell line (V9M) with untreated control and cyclophilin B as loading control. **f**, PLZF knockdown in normal cells. PLZF binding signal for control (untreated) and knockdown shown in purple, H3K27ac signal at sites bound by PLZF in control, and H3K27ac signal at sites with PLZF binding loss shown in blue. **g**, PLZF over-expression in CRC cells. PLZF binding signal for control (untreated) and over-expression shown in purple, H3K27ac signal at control and sites with PLZF binding gains shown in blue. **h**, Hierarchical clustering of H3K27ac RPKMs for reactivated H3K27ac sites (n=1636) in control and PLZF over-expression cells (OE-A, OE-B), PLZF bound L1s labeled in purple, top cluster annotated with L1 subfamily composition. **i**, CR TF binding overlap at PLZF responsive young L1s (L1HS, L1PA2, L1PA3 combined). CUT&RUN illustrates PLZF binding at young L1s with overexpression, representative of a normal/PLZF high context. CUT&RUN illustrates PLZF binding at young L1s in control cells, KLF5, and MYC, representative of a tumor/PLZF low context. **j**, Representative tracks of recurrently reactivated L1s in tumors sensitive to PLZF in CRC cell line, example track of recurrent samples for each young L1 subfamily. **k**, Model of L1 reactivation with PLZF and CR TFs.

To determine if PLZF regulates L1s in colon cancer, PLZF was knocked down in a normal colon cell line (FHC) and over-expressed in a colon cancer cell line low in PLZF activity (V9M) (Fig. 6e). CUT&RUN was then performed on treated cells for both PLZF and H3K27ac. In the normal cell line with high PLZF expression, PLZF knockdown leads to 73% of previously bound L1s losing PLZF binding at their 5’UTR with a coordinated increase in H3K27ac (Fig. 6f). Conversely, in the tumor cell line that has low PLZF, PLZF overexpression leads to 29% (476/1,636) of the reactivated fl-L1s gaining PLZF binding at their 5’UTR, with a large decrease in H3K27ac (Fig. 6g). Unsupervised clustering of H3K27ac RPKMs for all reactivated fl-L1s in the tumor cell that models L1 reactivation via PLZF loss (1636 total) and two replicates in which we rescue PLZF reveals a distinct cluster of activity changes in H3K27ac (Fig. 6h). L1s in the cluster are largely bound by PLZF when rescued and represent the elements that are most sensitive to reversing the effect of PLZF loss, and largely are sites with the highest H3K27ac signal. Reverting L1s are primarily composed of the three youngest L1 subfamilies (94%), which are also most associated with tumor-specific H3K27ac gains. We also find PLZF binds young L1s more frequently than older L1s when comparing all reactivated L1s prior to PLZF rescue, to those that are bound upon overexpression (Extended Data Fig. 8a-c).

Evaluating these PLZF responsive sites more closely for potential L1 activators upon PLZF loss, many CR TFs have binding overlap at these sites (Fig. 6i). At responsive L1s, we find both KLF5 and MYC present before PLZF rescue, both of which have been implicated in colon cancer progression^27–29^. We show in four recurrent L1s how H3K27ac signal in the tumor cell line with PLZF loss replicates the profiles of the tumor samples in our cohort almost perfectly. After PLZF rescue and binding at their 5’UTRs H3K27ac signal decreases dramatically to levels comparable or slightly above what we observe in the adjacent-normal samples. This demonstrates that individual L1 reactivation can be fully reversible with PLZF rescue (Fig. 6j). Together with the previous results, we propose a model in which PLZF loss leads to epigenetic derepression of L1s and exposes motifs for core regulatory factors, driving the oncogenic reactivation of young L1s (Fig. 6k).

## Discussion

We identify L1s as the family of REs most enriched in genomic elements that become active (gain H3K27ac) in the transition between adjacent-normal and tumor tissue in a cohort of colon cancer patients. L1 enrichment is especially strong in the elements that gain activity across multiple patients. The youngest L1s are the most likely to be recurrent. Generally, L1 activity was uniform across patients within subfamilies, varied widely between subfamilies, and increased gradually through the stages of tumor progression, consistent with previous models of hypomethylation. However, the three youngest L1 subfamilies deviated from the trend, exhibiting more rapid increases in activity and higher variability between patients. The degree of recurrence in young L1s also deviated from a random model of L1 activation, with many L1s active in the majority of patients. These observations suggest an additional mechanism of reactivation beyond the gradual increase in global hypomethylation driven by cell division.

Active L1s in the youngest subfamilies act as enhancers, gain loops to oncogenes and have pro-growth effects on cells. L1s are active in stem cells but remain poised in differentiated adult tissues (marked by H3K4me1). The 5’UTR of young L1s are enriched for motifs of core regulatory factors and we confirm binding of core factors at reactivated L1s. Furthermore, highly recurrent L1s contain a surplus of core factor motifs (MYC, KLF5, and others). However, the presence of motifs is insufficient for reactivation. We show that by modulating PLZF, a broad repressor of L1s, we can recapitulate tumor L1 reactivation in normal cells and subsequent core factor binding. Conversely, rescue of PLZF reverses L1 reactivation in tumor cells. The young L1 subfamilies are the most responsive to PLZF modulation. These observations support the existence of an additional, cell-division-independent mechanism of L1 reactivation, in which an abundance of core factor motifs in the consensus sequences of L1 5’UTR become exposed upon de-repression by loss of PLZF activity.

In contrast to the rare instances where an L1 disrupts a tumor suppressor or oncogene through insertional mutagenesis, oncogenic reactivation of young L1s is common across colon cancer patients. Importantly, alterations to PLZF and core factors that could initiate oncogenic L1 reactivation are also common. A survey of gains and losses in epigenetic and transcriptional activity of transcription factors in our cohort reveals that the most enriched core factors in young L1s are recurrently gained in tumor and that PLZF is recurrently lost in tumors. Because the excess activity in young L1s is highly frequent in patients, because individual L1s are recurrently active across virtually every patient, because of the ability of recurrently active L1s to act as enhancers to oncogenes, and because of the degree of wiring between the 5’UTRs of young L1s and the core regulatory circuitry of colon cancer, we propose oncogenic reactivation of young L1s as a hallmark of colon cancer. Our findings support the study of young L1s as early detection biomarkers, as has recently been suggested^30^, or in the treatment and prevention of cancer.

## Materials and Methods

### Cell Culture and tissue samples

A total of 46 colon tumors with paired adjacent-normals were acquired as surgically resected, flash frozen tissue in accordance with the Case Western Reserve University IRB protocols and approval. Across the cohort, 25 pairs were acquired from Cleveland,OH and 21 were acquired from Barcelona, Spain. Additional primary samples include 6 non-cancerous normal from patients with Crohn’s, Ulcerative Colitis, and Diverticulitis taken from non-diseased regions of the colon. Two adenomas from a patient with familial adenomatous polyposis profiled previously were also included in this work. Colon cancer cell line V9M, a cell line derived from colon tumors (cite) were cultured in MEM media (Gibco, cat. 10370-021) supplemented with 8% fetal bovine serum (Hyclone), 2mm L-glutamine (Gibco, cat. 25030-081), and 50 mg/ml 1 gentamycin (Gibco, cat. 15750-060). Normal colon line FHC (CRL-1831) was cultured in DMEM/F12 media (Gibco 11330-032) supplemented with 10% fetal bovine serum, 25mM HEPES, 0.005mg/ml insulin, 0.005mg/ml transferrin, 100ng/ml hydrocortisone, 20ng/ml human recombinant EGF, and 10ng/ml cholera toxin.

### ChIP-seq

ChIP-seq was done as previously described^22, 31^ using an antibody targeting H3K27ac (Abcam, ab4729), and H3K4me1 (Abcam, ab8895). In brief, frozen tissue was crushed to a powder in 4°C and crosslinked with methanol free formaldehyde following Covaris truChIP Chromatin Shearing Kit protocol. Libraries were prepared as previously described and sequenced at CWRU Genomics Core Facility and NYGC.

### ChIP-Seq and CUT&RUN data processing

Cutadapt v1.9.1^32^ was used to remove paired-end adapter sequences and discard reads with a length less than 20bp. All FASTQs were aligned to both the hg19 genome assembly (retrieved from hgdownload.cse.ucsc.edu/goldenPath/hg19/chromosomes) and the *E. coli* K12 MG1655 genome assembly (retrieved from igenomes.illumina.com.s3-website-us-east-1.amazonaws.com/Escherichia_coli_K_12_MG1655/NCBI/2001-10-15) using BWA-MEM v0.7.17-r1188 with default parameters in paired-end mode. Output SAM files were converted to binary (BAM) format, sorted, indexed, and PCR duplicates were removed using SAMtools v1.10. Peaks were detected for hg19-aligned samples with MACS v2.1.2^33^ using default parameters for transcription factors or with the ––broad flag set for histone marks and both types of marks utilized –format=BAMPE. SAMtools v1.10 was used to determine the number of uniquely mapped *E. coli* spike-in reads. Spike-in normalization factors were computed by taking the reciprocal of the number of uniquely mapped *E. coli* reads divided by the total number of reads after removal of PCR duplicates such that after normalization the *E. coli* spike-in signal is set to be equal across samples. DeepTools v3.2.0^34^ was used to generate spike-in-normalized bigWig tracks with 50 bp bin sizes from the final sample BAM files for the hg19 alignments using the –scaleFactor option set according to the appropriate spike-in normalization factor for a given sample. BigWigs were visualized on the Integrative Genomics Viewer^35^ in order to eliminate samples with pronounced track irregularities or low signal-to-noise ratio.

### RNA-seq

A subset of tissue was crushed and saved in 1mL trizol at –80 for RNA-seq, RNA was extracted with trizol following invitrogen RNA isolation protocol. The quantity and quality of RNA were evaluated using both Nanodrop (ThermoFisher Scientific) and Bioanalyzer 2100 (Agilent Technologies). RNA-seq libraries were prepared and sequenced at CWRU Genomics Core Facility.

### RE enrichment survey

We intersect the tumor exclusive gained peaks and 100 null peak sets of enhancer elements sampled from the source H3K27ac peaks at random with 1,346 RE subfamilies (excluding simple repeats). Z-scores derived from the null sets reveal gVELs are primarily depleted of REs. RE families SINE MIR, Low complexity, SINE Alu and LINE L2 make up the bottom ten, while LINE L1, Satellite centr, LTR ERVL-MaLR and LTR ERVK make up the top ten.

### H3K27ac overlap test

All H3K27ac peaks were intersected with 8,679 full-length LINE1 elements annotated by RepeatMasker (https://www.repeatmasker.org). We restricted our analysis to full-length L1’s, as these elements retain the 5’ UTR that contains regulatory elements. For each recurrence level and subfamily pair we sample approximately the same number of reactivating gained VEL. We use 50 permutations for the overview scan across all subfamilies and recurrence levels and fit a negative binomial distribution to the null and compute an exact p-value. We ran the H3K27 overlap test one-sided and tested 26 subfamilies × 36 recurrence levels (936 tests). The *p*-value reported in the main text is corrected using the Bonferroni method.

### L1 isolation analysis

The effect of regulatory elements neighboring L1s could possibly inflate the reactivation rates we calculated. In an effort to overcome this we set out to estimate the “intrinsic” reactivation rates of each subfamily, that is, the fraction of the reactivation rate that is driven by the sequence of each L1 instance and not neighboring regulatory elements. We found that, although L1s are more likely to reactivate when surrounded by other active regulatory elements, the trend of accelerating activity in the young L1 subfamilies is maintained. We corrected this “sympathetic” by recalculating reactivation rates in isolated L1 instances with increasingly large flanking regions with no active elements (10 to 50 kb). Intrinsic reactivation rates appear to plateau between 25 and 50 kb of isolation, confirming a sequence-driven mechanism of reactivation.

### L1 subfamily recurrence test

We devised a score to measure the degree of recurrence or non-random patterning by shuffling activity states across a binary matrix of L1 instances versus samples, using a single reactivation rate for each subfamily and patient combination (fig. S3D). This process allows us to establish an expected number of highly recurrent L1 instances under a model of fully random reactivation. By measuring the relative enrichment of observed over expected recurrence we can score the degree of deviation from randomness for each of the three cell states and each of the 27 subfamilies (81 tests total). We quantify the difference between observed and expected reactivation patterns using Levene’s test for homogeneity of variance across groups from the “car” R package. We found that although most L1 subfamilies exhibit patterns of reactivation that are consistent with a random model, the youngest L1 subfamilies showed a markedly non-random pattern.

### Motif analyses

TF motif enrichment analyses were performed with HOMER software using the findMotifsGenome tool (http://homer.salk.edu/homer/index.html). The 5’UTR of L1s with H3K27ac signal were motif mined with standard parameters. FIMO from the MEME suite tool was used to generate bed files for enriched TFs to calculate the fraction of overlap for each L1 subfamily^36^. Hierarchical clustering and heatmaps of overlaps were generated with Morpheus (https://software.broadinstitute.org/morpheus). Additional colocalization analyses were performed with the enrichment analysis tool in the ChIP-Atlas database (https://chip-atlas.org). A total of 1459 digestive tract specific TF ChIP-seqs were intersected with regions of interest using a randomized background (permutation x100). In all cases, the most significant TF was reported in cases of TF redundancy within the output. To identify motifs specifically enriched at recurrent L1s, a set of 326 isolated L1PA2s were scanned for core factor motifs, comparing instances that do not reactivate to each of the clusters showing varying degrees of recurrence.

### Core Regulatory Circuitry

To identify core regulatory circuitry Colton (https://github.com/linlabcode/CRC) was used where ROSE super-enhancer associated TFs are used to calculate inward and outward degrees of connectivity. For expression, a TPM cutoff of 3 was used for a given TF to be considered within the CRC. A TFs inward and outward regulation was determined, how often it binds its own SE and the SEs of others respectively. Taken together each TF will have a total degree score to reflect the degree of connectivity within the circuit.

### L1 TAD Gene Expression Analysis

To determine the regulatory potential of active fl-L1s, we used all recurrent L1s as well as case-specific instances to compare the expression of target genes in patients with vs patients without the reactivated elements. In total, 19,194 L1 peak x gene combinations were tested (median of 11 genes per L1 peak). Our analysis found 432 unique genes whose expression correlated with at least one of 291 reactivated L1s. To address the possibility that these associations mainly reflect genes that are over-expressed in cancer, with L1 reactivation being only a correlate of tumor status, we regressed gene expression on L1 peak and tumor status of the sample. For the majority of correlating L1 X gene pairs (334/497, or 67%), the presence of the L1 peak, either alone or conditioned on tumor status, was a significant predictor of gene expression.

### HiChIP sample preparation

HiChIP was performed as previously described with the Dovetail MNase-HiChIP kit6 (Dovetail, 21007) with a few modifications. For each experiment, 10×106 cells were collected and snap-frozen at –80C for a minimum 30 minutes then cross-linked with disuccinimidyl glutarate (DSG, Thermo Fisher, A35392) and 37% formaldehyde (Sigma-Aldrich, F8775).

### HiChIP analysis

Paired-end reads were mapped to the human genome (build hg38) and mouse genome (build mm10) to align human genomic content and mouse spike-in controls respectively, using Bowtie2 within the HiC-Pro pipeline (PMID: 26619908). Mapped reads were then filtered for junction validity, and read pairs with contact range of 1000bp or less were removed to filter out spurious MNase ligation products. PCR duplicate-read pairs of punctate HiChIP pull-downs such as AR and CTCF were not removed since they piled up at ChIP-seq peaks rather than acting as randomly distributed noise. Final read pairs (allValidPairs) were converted to Juicebox compatible.hic files for visual inspection using the hicpro2juicebox script provided within HiC-Pro (PMID: 33743768). All downstream analyses requiring sub-matrix extraction from.hic files were performed using Juicer and strawr (PMID: 27467249).

### APA plots

We used APA (Aggregated Peak Analysis) plots to visualize 3D contexts. APA is used to show whether (differential) loop signals are isolated from their surrounding contexts and amplify the signal after collapsing 3D loops at their center. APA was implemented using Juicer tools (v1.19.02, https://github.com/aidenlab/juicer) and custom R scripts. We ran the apa module with no internal normalization and no minimum distance from the diagonal threshold at 10kb resolution.

### CUT&RUN

TF and histone marks were profiled according to Epicypher CUTANA ChIC / CUT&RUN Kit V3 (14-1046). Briefly, 500,000 cells were harvested in PBS, washed and incubated overnight with target antibodies at 4°C, 0.5ug for histone marks (H3K27ac, H3K4me3) and 1ug for TFs (ETV4, CDX2, TCF7L2, KLF4, KFL5, MYC, and PLZF). Library prep was performed with NEBNext Ultra II DNA Library Prep Kit for Illumina per the manufacturer’s instructions. For histones, 1X AMPure beads were used for cleanup and size selection of fragments. For TFs, 1.75X AMPure beads were used for cleanup and size selection to retain smaller fragments (<120bp). Bioanalyzer and qubit were used to quantify and QC final libraries. Libraries were sequenced on the NovaSeq at Novogene (Sacremento, CA).

### Knockdown of PLZF

Horizon Discovery ON-TARGETplus Human ZBTB16 SMARTpool siRNAs (4 siRNA/gene target, L-018719-00-0005) were used to transfect FHC cells with PLZF targeting siRNA (25nM) following manufacturer’s protocol. Non-targeting siRNA (D-001810-01-05) and siGLO red transfection control (D-001610-01-05) were used to monitor off-target toxicity and transfection efficiency. 25,000 cells were plated into 12 well plates and were transfected for 48hrs. Cells were collected at 72hrs for downstream expression analyses and CUT&RUN.

### Overexpression of PLZF

PLZF was overexpressed using TetO-FUW-pgk-puro, which was a gift from Emily Dykhuizen (Addgene plasmid # 85747). Lentivirus containing the overexpression plasmid was generated using LentiX Packaging Single Shots (Clontech, 631278) according to the manufacturer’s protocol. V9M cells were then transduced with the PLZF lentivirus by spinfection. In brief, 5×10^4 cells were seeded in 12-well plates and allowed to adhere for 24 hours. Media was then aspirated and replaced with 0.3 mL of unconcentrated lentiviral supernatant, 0.7 mL of fresh media, and polybrene to a final concentration of either 8 ug/mL (condition 1) or 4 ug/mL (condition 5). Plates were then spun at RT for 30 minutes at 800 x g. Once plates were done spinning, they were placed in a 37°C incubator overnight. Lentivirus containing media was aspirated the next day and replaced with media containing 1 ug/mL doxycycline hyclate (Cayman, 14422). Cells were then grown for an additional 48 hours before being collected for western blot and CUT&RUN analysis.

### Western Blot

Cells were lysed with RIPA buffer containing 1X protease inhibitor. (Roche, 4693159001). Protein was isolated and concentration was measured utilizing a BCA assay kit (Thermo Fisher, 23225). 12.5ug of protein was loaded onto precast 4%–12% Bis-Tris gels (Invitrogen, NP0321BOX) and run at 200V for 35 minutes. Protein was then transferred to PVDF membranes (Bio-Rad, 1704157) using the trans-blot turbo transfer system (Bio–Rad, 1704150). The membrane was then blocked with dry milk that was diluted to 5% using PBST for 1 hour at room temperature. After the block, membranes were incubated overnight at 4℃ with primary antibodies PLZF (MA5-15667) and cyclophilin B (abcam, catalog ab16045). Membranes were then washed with PBST and incubated with HRP-conjugated secondary antibodies (Santa Cruz Biotechnology Inc., catalog sc-2004; Cell Signaling Technology, catalog 7076S) at room temperature for 1 hour. Chemiluminescence was then conducted with SuperSignal West Pico Plus Chemiluminscent Reagent (Thermo Fisher, X8340773) and membranes were imaged using ChemiDoc Touch Imaging system (BioRad, 1708370).

---

**Extended Data Fig. 1.**
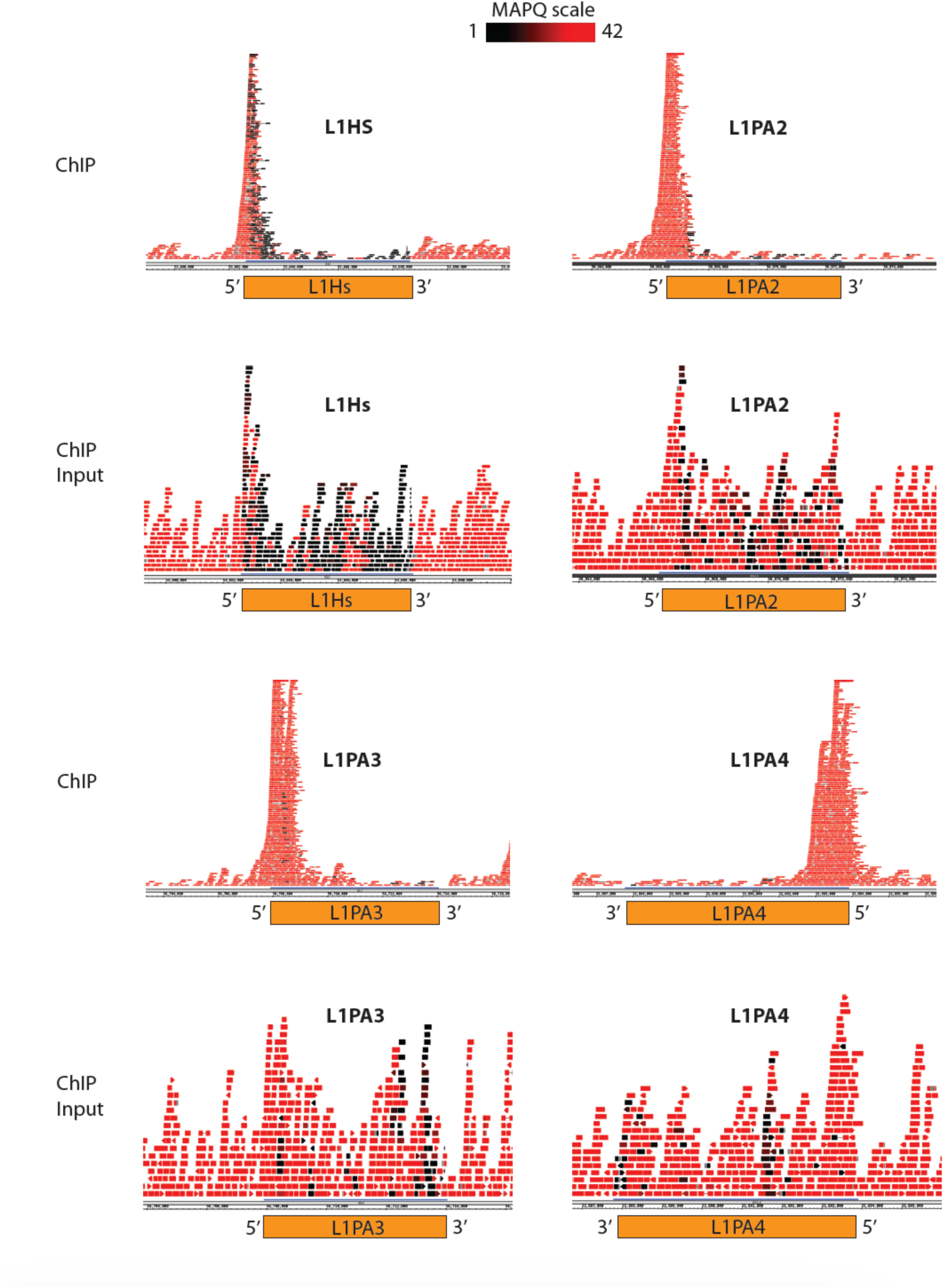
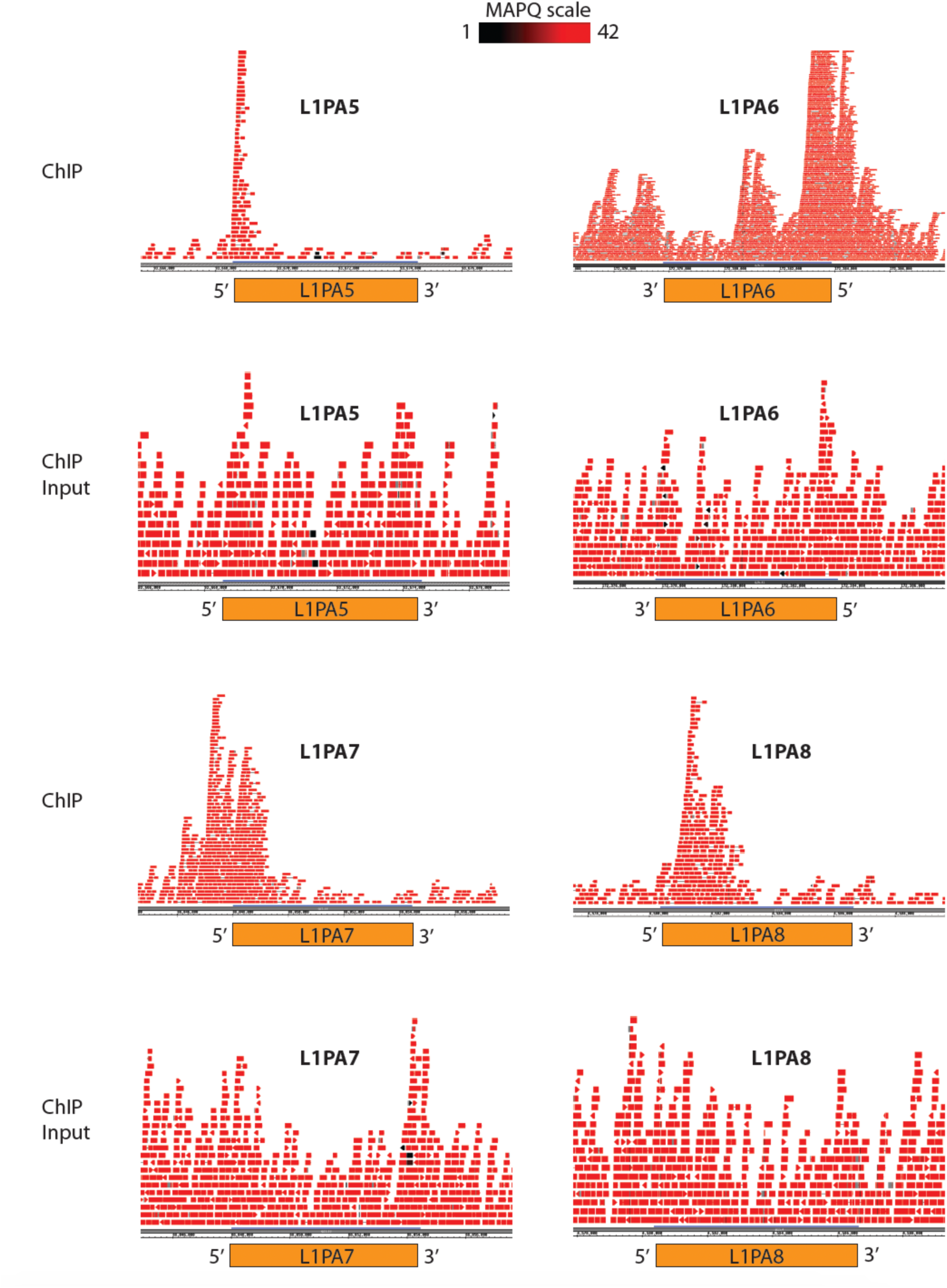
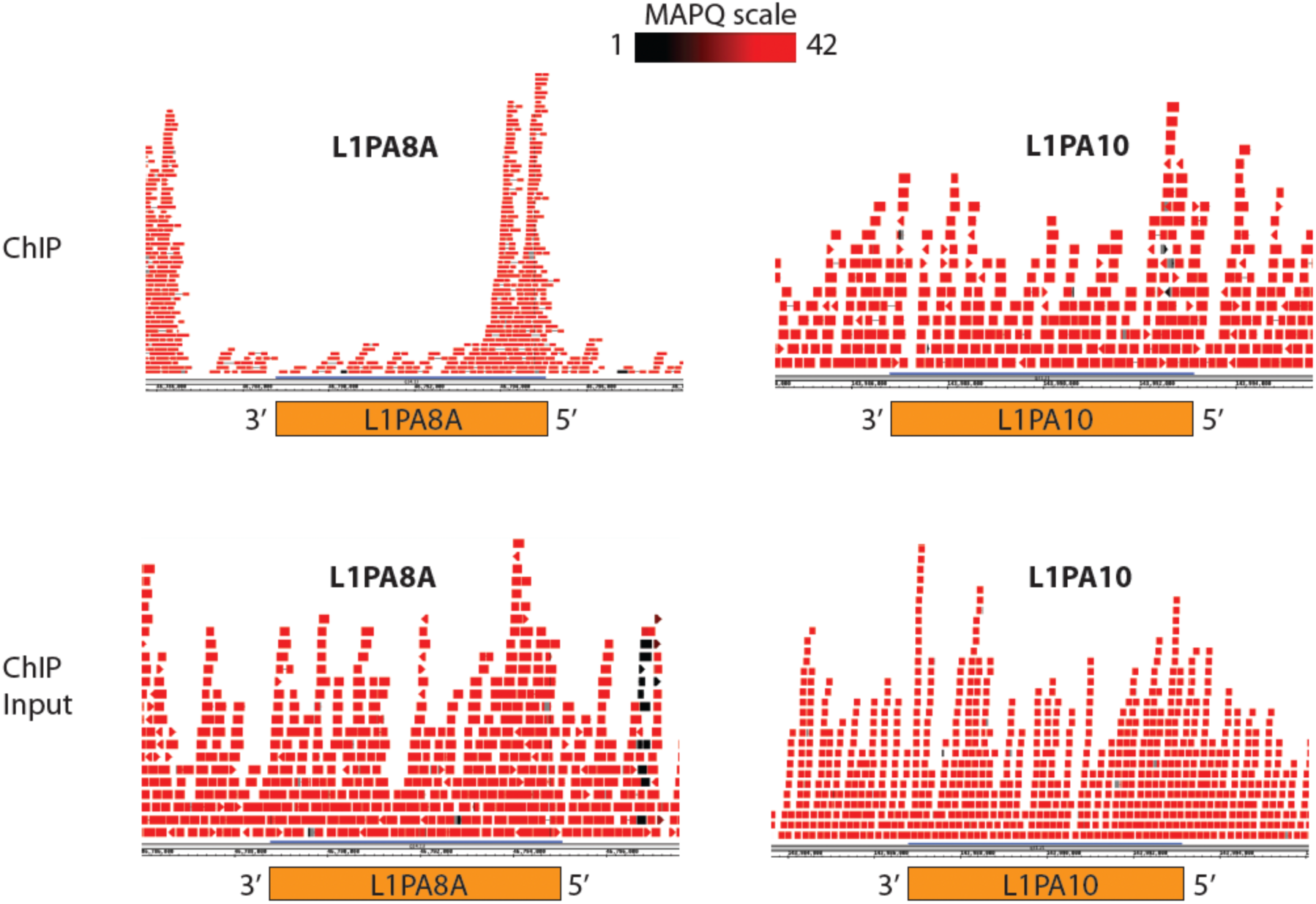
fl-L1 Mappability with ChIP-seq. ChIP–seq reads mapped to individual L1 instances in the genome. The color scale indicates the mapping quality score MAPQ for each read pair. MAPQ = 10log10p, where p is the probability that true alignment belongs elsewhere. The bodies of L1 repeats are uniquely mappable and exhibit distinct pileup in the 5’ region of the examined L1 subclasses. Examples of mapping are shown for both input and H3K27ac ChIP-seq.

**Extended Data Fig. 2.**
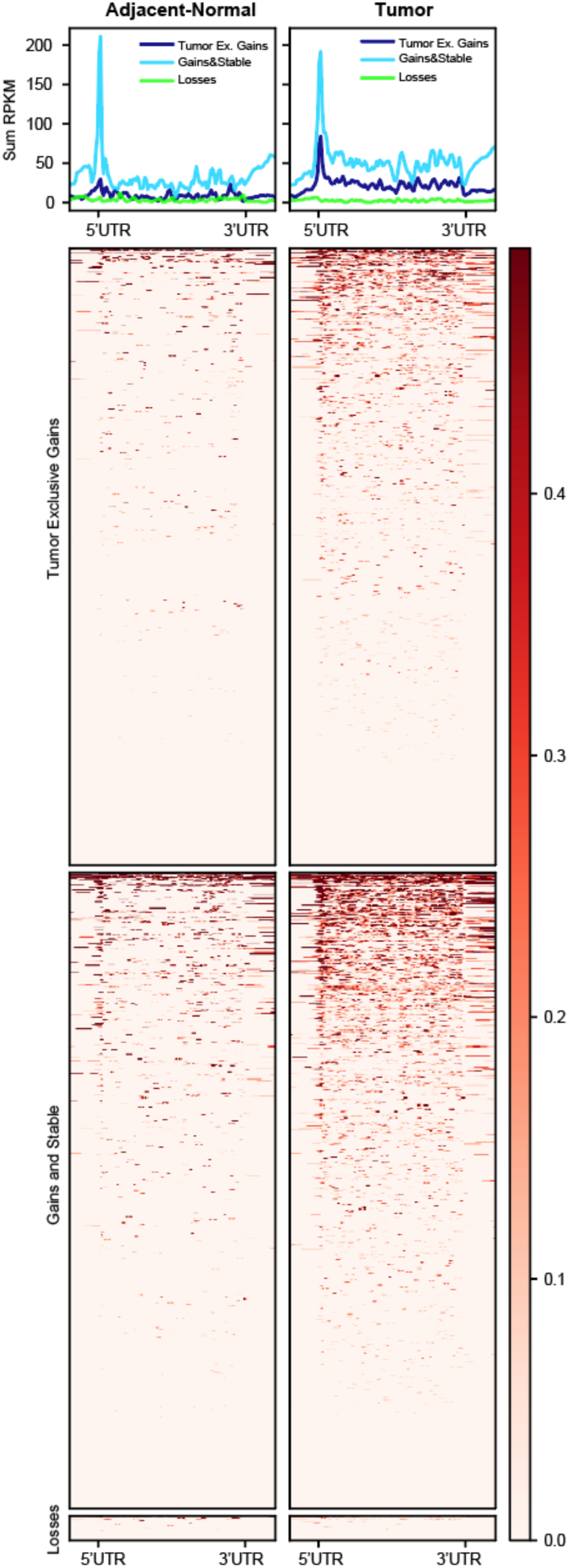
fl-L1 tumor gains show increased transcription. RNA-seq RPKM signal at fl-L1 for tumor exclusive gains, gains, stables, and losses.

**Extended Data Fig. 3.**
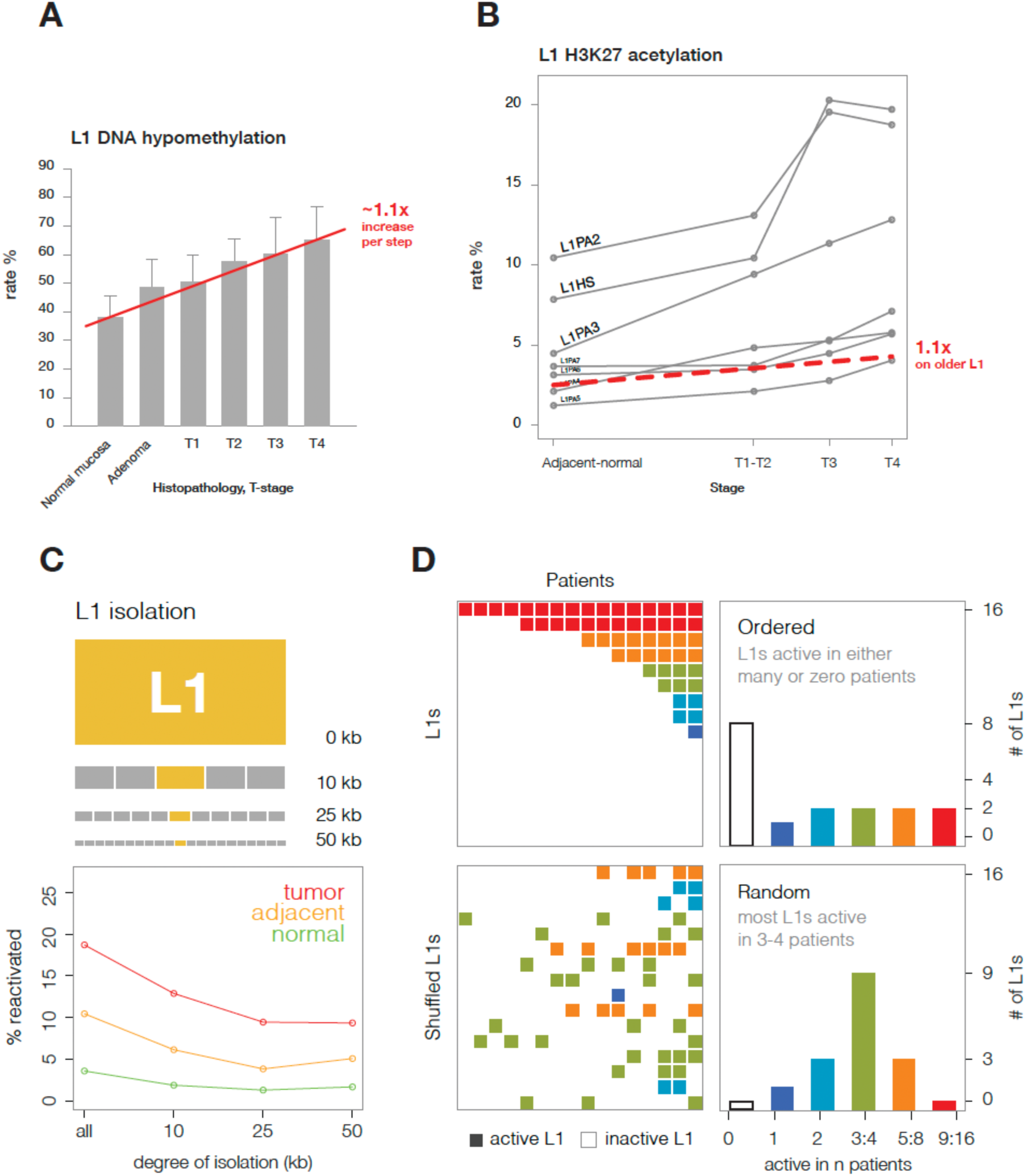
Young L1s have greater activity and recurrence than expected. **a**, Rate of L1 DNA hypomethylation (y-axis, %) across tumor progression (adapted from Sunami et al. 2011). Average increase per step in red. **b**, Rate of L1 H3K27ac acetylation (y-axis, %) across tumor progression separated by subfamily. Expected increase in older L1 subfamilies based on DNA hypomethylation progression in read. **c**, L1 isolation diagram (top). Percentage of reactivated L1PA2s (y-axis) versus size of flanking regions used to isolate the L1s (x-axis, bottom). **d**, Illustration of random and non-random (or highly recurrent) L1 reactivation patterns (left). Number of instances (y-axis) versus binned recurrence levels (x-axis, right)

**Extended Data Fig. 4.**
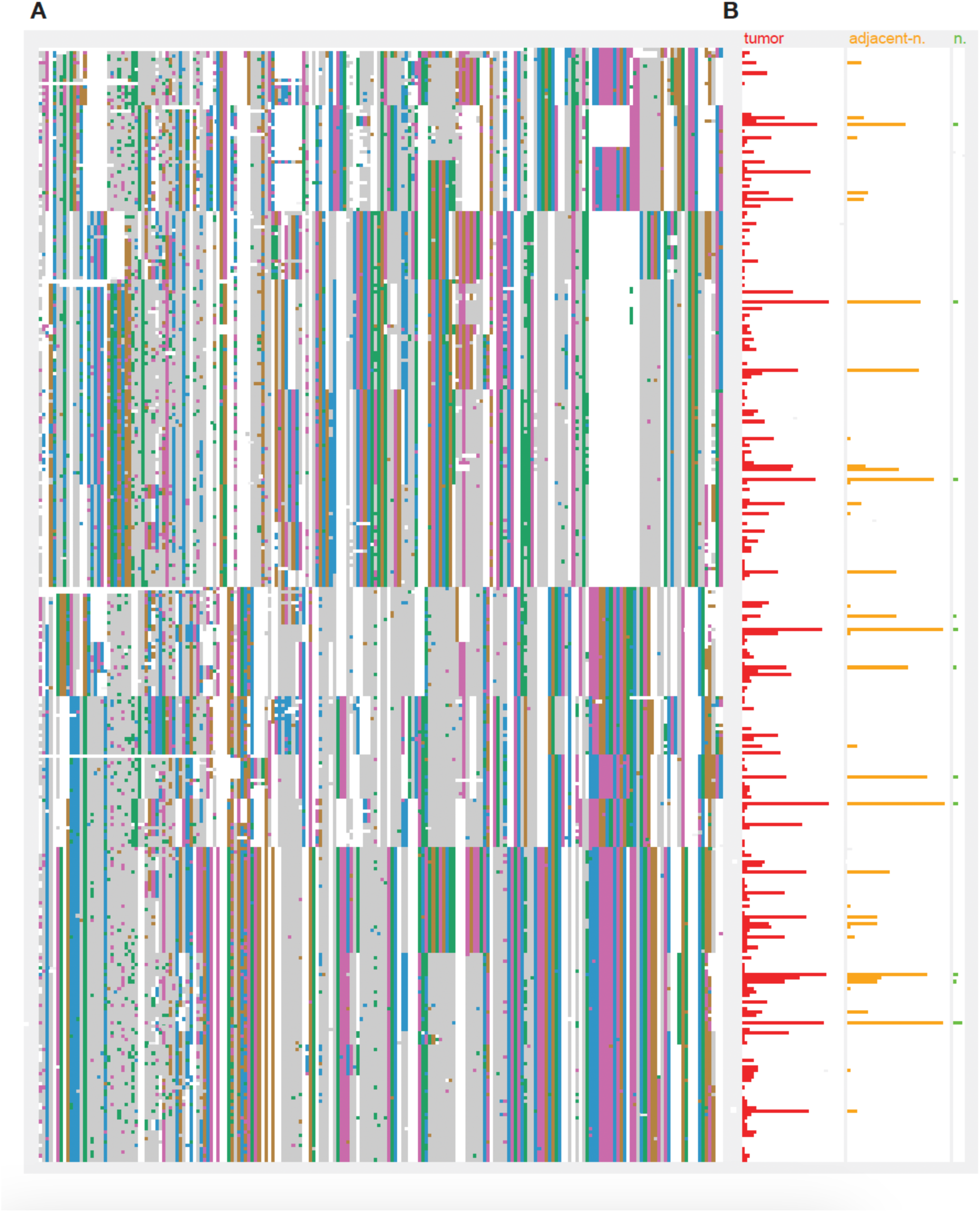
L1PA2 sequence alignment for high entropy bases and recurrence in cohort. **a**, Alignment of all full-length L1PA2s clustered based on sequence similarity. Sequence variants are filtered with entropy to prioritize base distributions favorable to splitting each subfamily into large subgroups. **b**, Recurrence levels for tumor (red), adjacent-normal (yellow), and non-cancerous normal (green).

**Extended Data Fig. 5.**
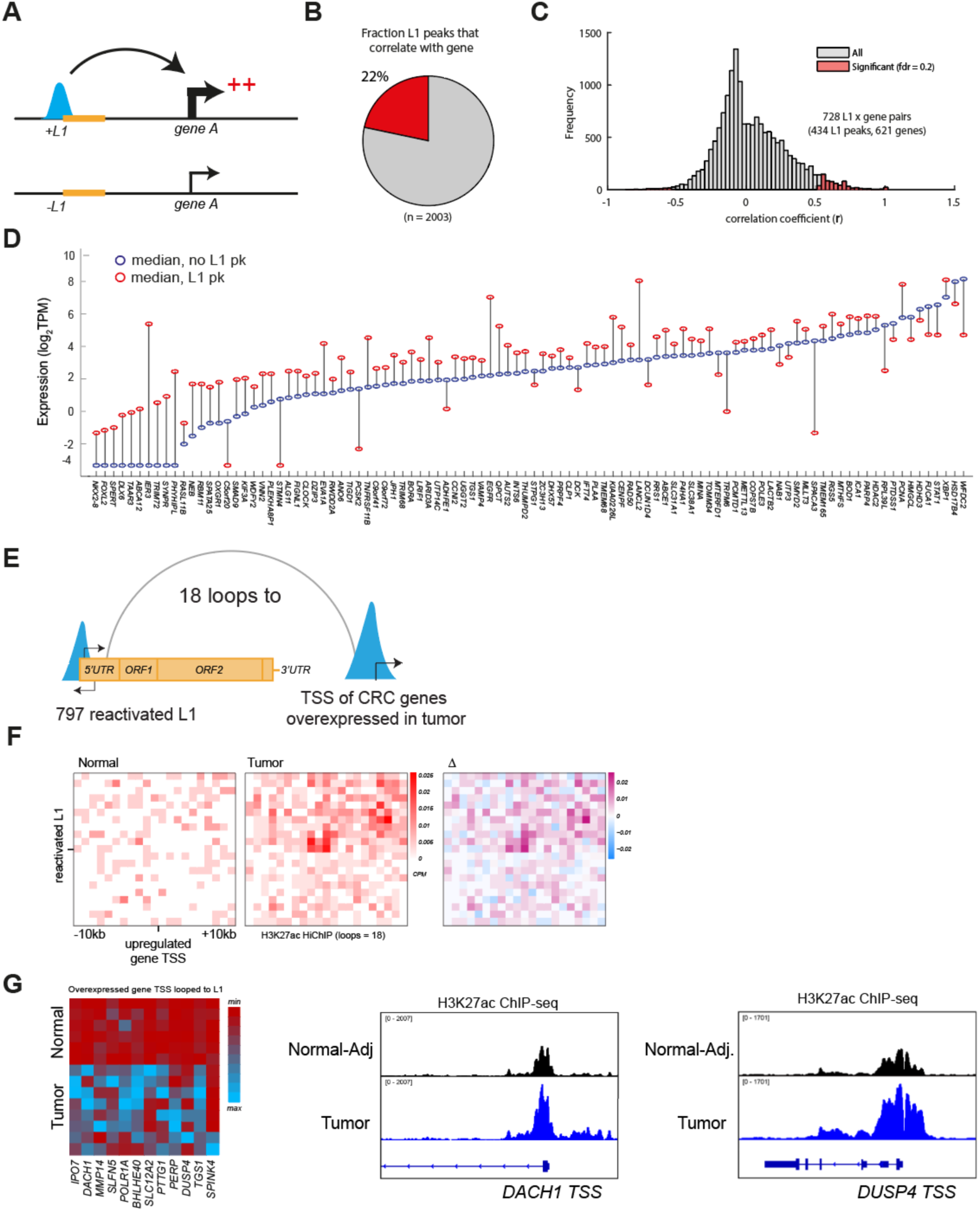
L1s correlate with aberrant expression of nearby genes and loop to upregulated genes in tumors. **a**, Schematic of L1 enhancers-gene correlation analysis. The presence or absence of L1 enhancers was correlated with expression of all genes within the same topologically associated domain (TAD). **b**, Fraction of L1 enhancers with significant correlation with expression of nearby gene. **c**, Distribution of correlation coefficients for all L1 enhancer-gene pairs. Significant interactions are highlighted in red. **d**, Top 100 putative target genes of L1, sorted by correlation coefficient. Blue and red circles represent median expression of samples lacking L1 enhancer and harboring L1 enhancer, respectively. **e**, Summary of HiChIP results, reactivated L1s in sample 28T (n=797) form new loops to 18 TSS of overexpressed genes (n=238) in colon tumors. **f**, Contact frequencies of L1-TSS loops, defined by reactivated L1s and TSS within 5 MB and annotated based on CPM from H3K27ac HiChIP in tumor. **g**, Expression levels in normal and tumor samples across genes overexpressed from TSS looped to L1s in tumor. H3K27ac ChIP-seq tracks for example genes *DACH1* and *DUSP4* with reads showing upregulated TSS in tumor compared to normal.

**Extended Data Fig. 6.**
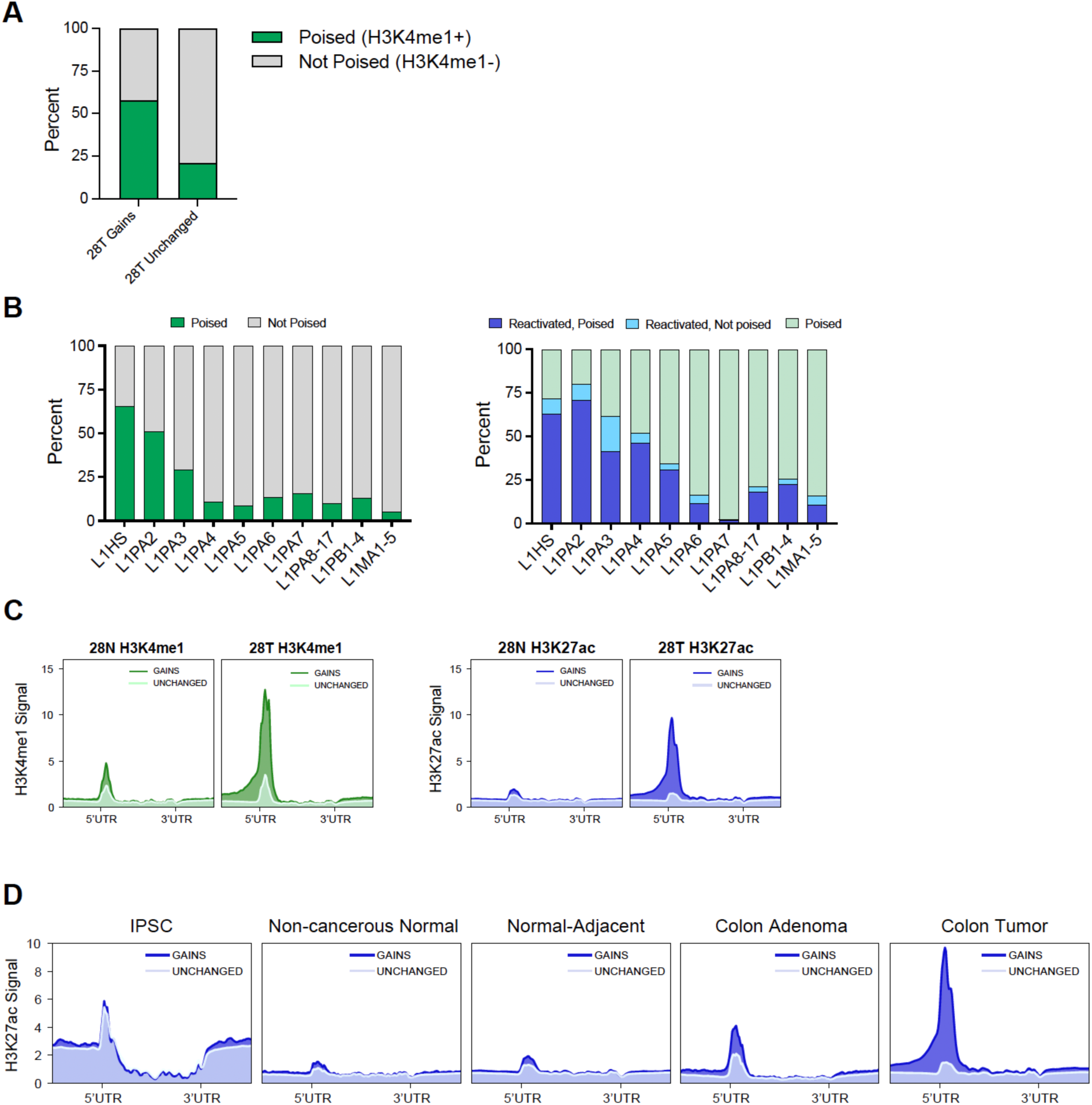
Poised and L1 reactivation data for sample 28. **a**, Experimental outline for H3K4me1 ChIP-seq (CRC-14 & CRC-28), H3K4me1 sites intersected with sample specific H3K27ac gains to calculate the percent of L1s gains poised in adjacent normal. **b**, Percent of L1 subfamilies poised in normal (green), percent poised L1s that become reactivated in tumor (blue), percent of L1s not poised in normal but are reactivated in tumor (light blue). **c**, H3K4me1 and H3K27ac ChIP-seq signal at gains compared to unchanged fl-L1s. **d**, H3K27ac signal at gains and unchanged fl-L1s across different stages and cell types illustrating developmental progression of L1 reactivation.

**Extended Data Fig. 7.**
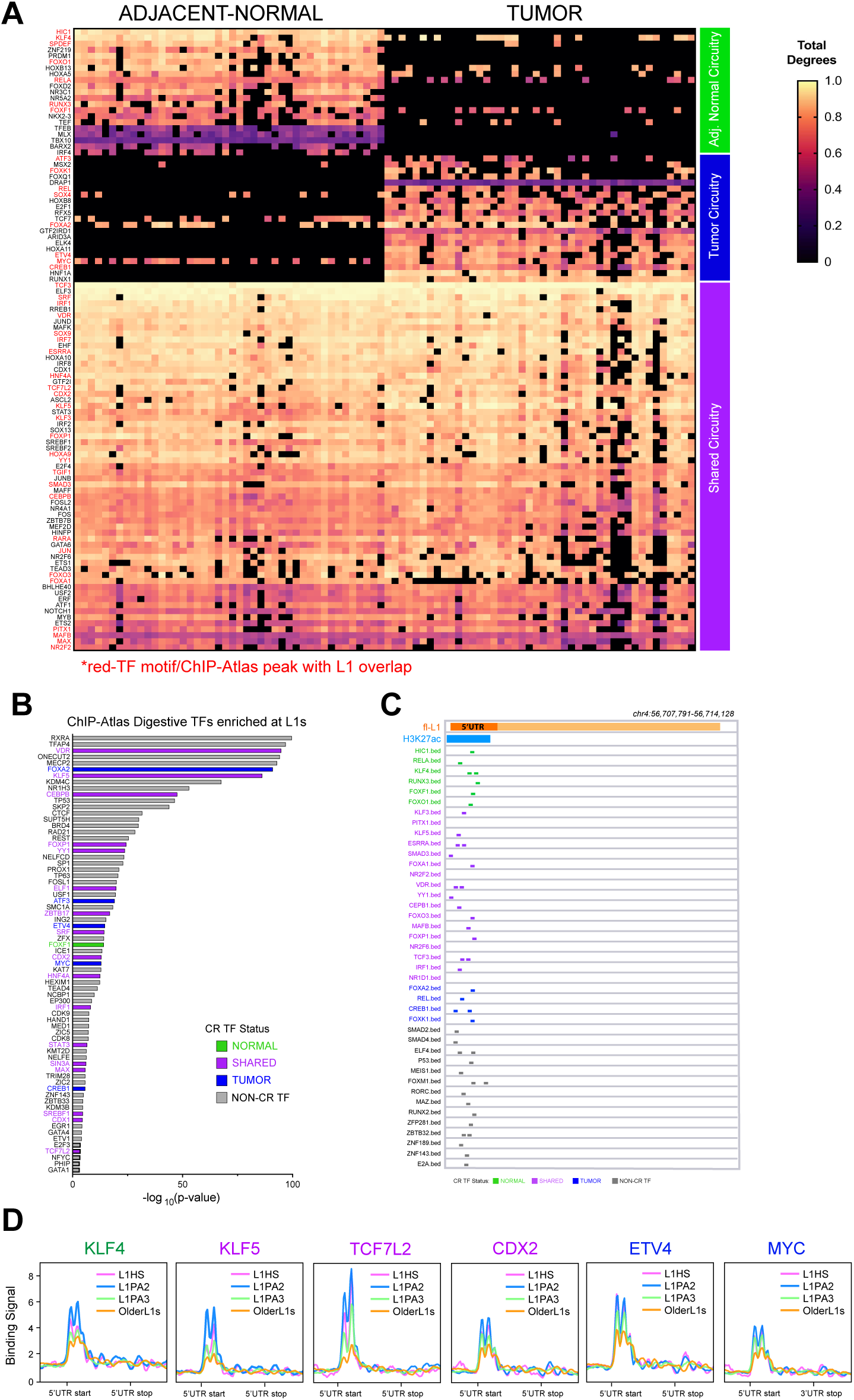
Additional core circuitry data, ChIP-atlas enrichment, and motif locations. **a**, Heatmap of top 100 core regulatory circuitry TFs, TFs enriched at L1s annotated in red. Heatmap values represent the total degrees of connectivity. **b**, TF binding overlap (ChIP-atlas database) for reactivated L1s. **c**, motif distribution example within the 5’UTR of fl-L1, CR TF status color labeled. **d**, CR TF binding signal at the 3 youngest subfamilies and combined older L1 subfamilies at the L1 5’UTR.

**Extended Data Fig. 8.**
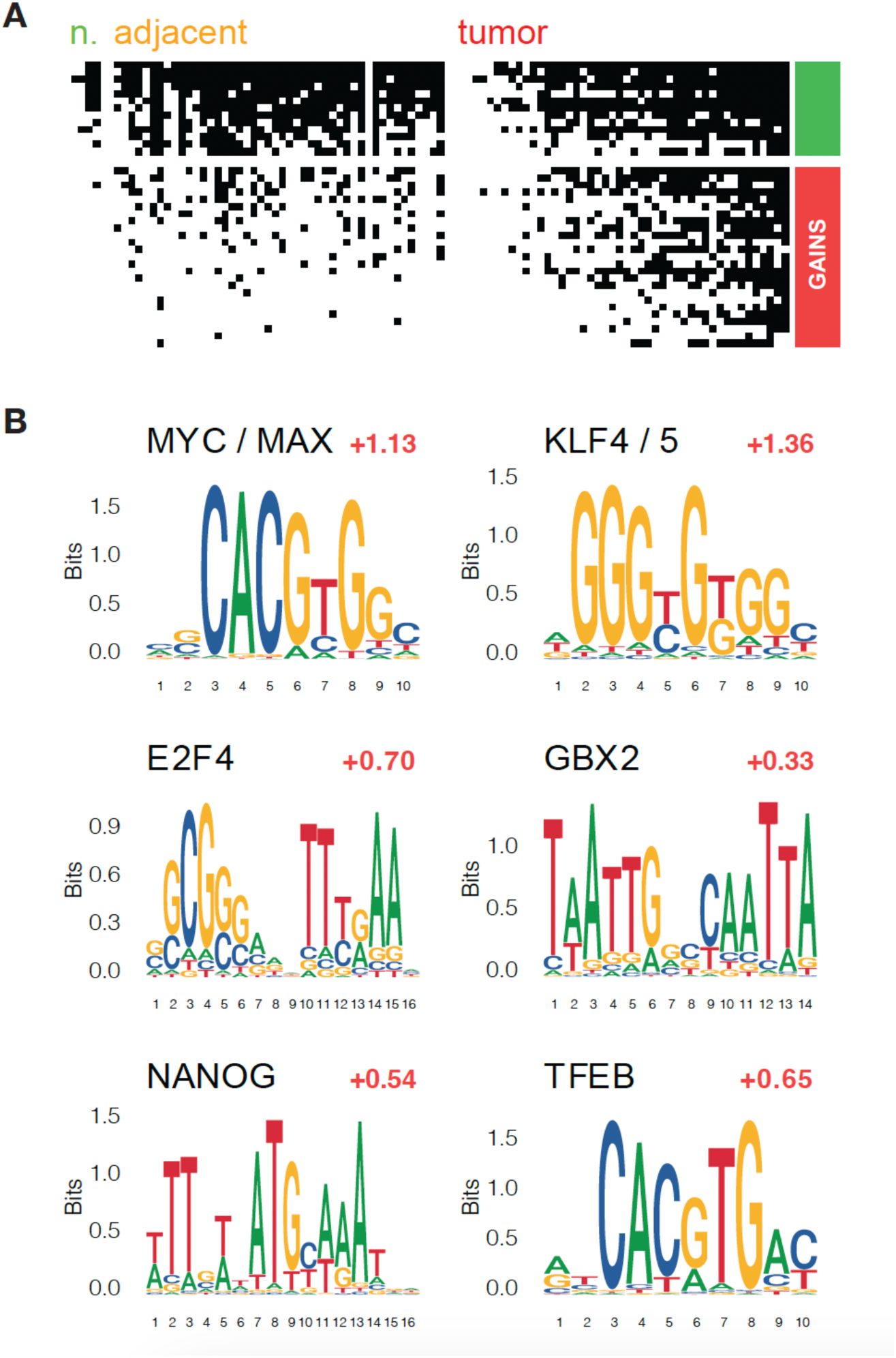
L1 motif enrichment in recurrent L1PA2. **a**, Binary heatmap of TF motif enrichment at recurrent L1PA2s compared to L1PA2s without H3K27ac signal in non-cancerous normals, adjacent-normals, and tumors. Top category (green) represents motifs shared across categories, bottom (red) represents TF motifs gained in tumors. **b**, Motif logos for L1PA2 instances in the tumor cluster with the highest recurrence contain on average an additional motif for both MYC/MAX and KLF4/5, other top CR TF enrichments for E2F4, NANOG, GBX2, TFEB.

**Extended Data Fig. 9.**
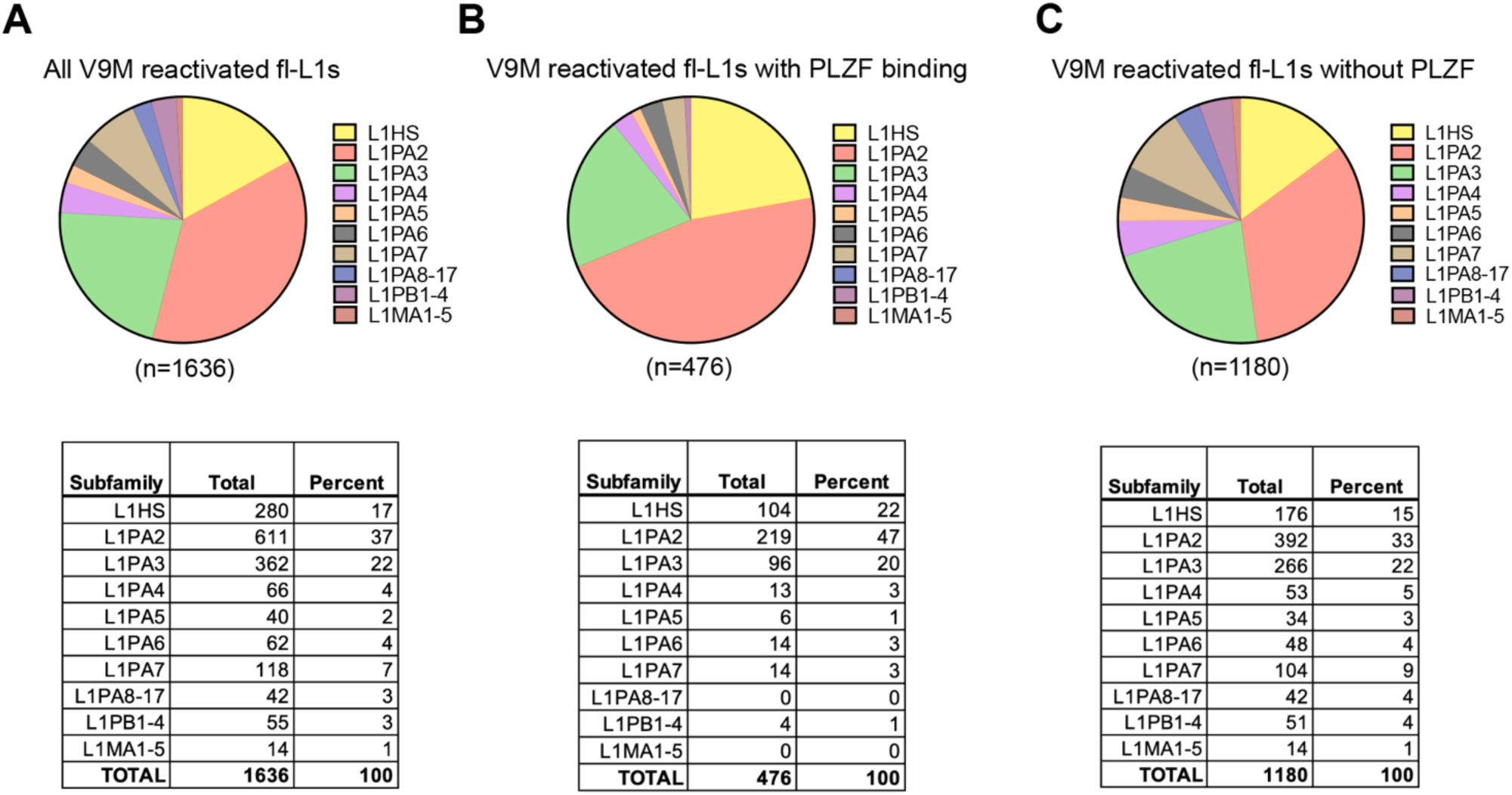
PLZF binding at L1 subfamilies. **a**, Percent of all reactivated L1s (n=1636) in control line V9M broken down by subfamily **b**, Percent of reactivated L1s bound by PLZF upon overexpression (n=476) broken down by subfamily. **c**, Percent of reactivated L1s not bound by PLZF upon overexpression (n=1180) broken down by subfamily. Tables display total counts and percentages shown in pie charts.

**Supplementary Table 1.**
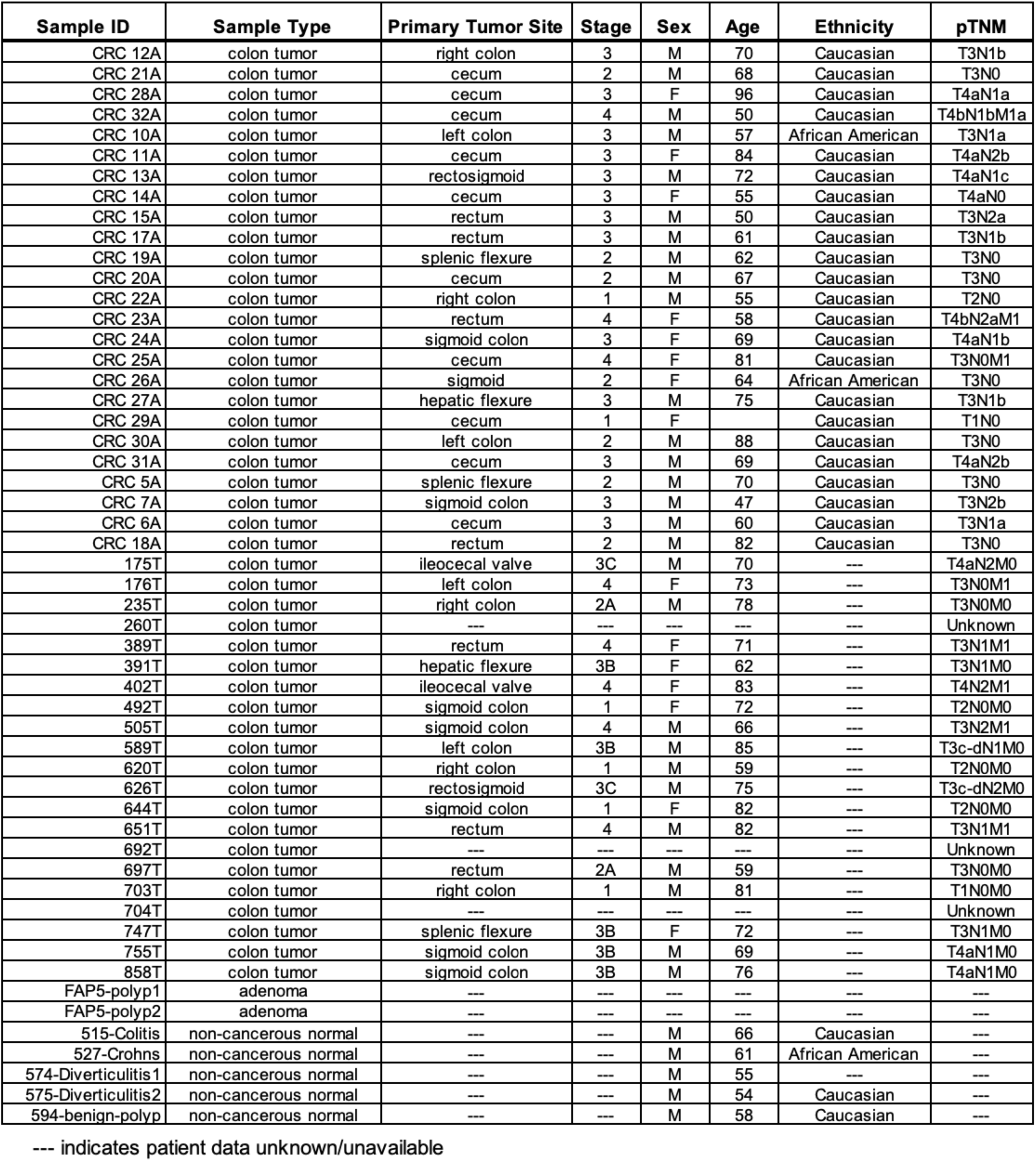
Clinical features of colon samples profiled in this study.

**Supplementary Table 2.** Characteristics of L1 subfamilies including totals for all length (truncated L1s and full length identified with RepeatMasker), totals for only full length L1s. Totals also shown for tumor exclusive gains only (Tx). Fraction of total indicates full length L1s / all length L1s for each subfamily.

| subfam | all_len_total | all_len_Tx | full_len_total | full_len_Tx | frac_total | frac_Tx | expected_fl_Tx | fold_change |
| --- | --- | --- | --- | --- | --- | --- | --- | --- |
| L1HS | 1571 | 229 | 314 | 184 | 19.987 | 80.349 | 46 | 4 |
| L1PA2 | 4918 | 500 | 1039 | 437 | 21.126 | 87.4 | 106 | 4.1 |
| L1PA3 | 10703 | 650 | 1488 | 531 | 13.903 | 81.692 | 90 | 5.9 |
| L1PA4 | 11926 | 470 | 1340 | 306 | 11.236 | 65.106 | 53 | 5.8 |
| L1PA5 | 11398 | 312 | 998 | 148 | 8.756 | 47.436 | 27 | 5.5 |
| L1PA6 | 6051 | 203 | 806 | 105 | 13.32 | 51.724 | 27 | 3.9 |
| L1PA7 | 13116 | 327 | 930 | 106 | 7.091 | 32.416 | 23 | 4.6 |
| L1PA8 | 8111 | 134 | 200 | 13 | 2.466 | 9.701 | 3 | 4.3 |
| L1PA8A | 2429 | 45 | 124 | 7 | 5.105 | 15.556 | 2 | 3.5 |
| L1PA10 | 7090 | 152 | 109 | 4 | 1.537 | 2.632 | 2 | 2 |
| L1PA11 | 4144 | 75 | 69 | 7 | 1.665 | 9.333 | 1 | 7 |
| L1PA12 | 1769 | 42 | 11 | 0 | 0.622 | 0 | 0 | NaN |
| L1PA13 | 9060 | 132 | 86 | 1 | 0.949 | 0.758 | 1 | 1 |
| L1PA14 | 3056 | 54 | 29 | 0 | 0.949 | 0 | 1 | 0 |
| L1PA15 | 8463 | 137 | 77 | 3 | 0.91 | 2.19 | 1 | 3 |
| L1PA16 | 13973 | 255 | 69 | 6 | 0.494 | 2.353 | 1 | 6 |
| L1PA17 | 4789 | 90 | 16 | 1 | 0.334 | 1.111 | 0 | Inf |
| L1PB1 | 13093 | 360 | 292 | 44 | 2.23 | 12.222 | 8 | 5.5 |
| L1PB2 | 2887 | 55 | 30 | 2 | 1.039 | 3.636 | 1 | 2 |
| L1PB3 | 3601 | 64 | 22 | 3 | 0.611 | 4.688 | 0 | Inf |
| L1PB4 | 7440 | 112 | 19 | 1 | 0.255 | 0.893 | 0 | Inf |
| L1MA1 | 4314 | 58 | 71 | 2 | 1.646 | 3.448 | 1 | 2 |
| L1MA2 | 7417 | 108 | 96 | 5 | 1.294 | 4.63 | 1 | 5 |
| L1MA3 | 8888 | 147 | 96 | 5 | 1.08 | 3.401 | 2 | 2.5 |
| L1MA4 | 10000 | 131 | 33 | 1 | 0.33 | 0.763 | 0 | Inf |
| L1MA5 | 4447 | 84 | 15 | 0 | 0.337 | 0 | 0 | NaN |

